# A parasitoid wasp of *Drosophila* employs preemptive and reactive strategies to deplete its host’s blood cells

**DOI:** 10.1101/2021.04.30.442188

**Authors:** Johnny R. Ramroop, Mary Ellen Heavner, Zubaidul H. Razzak, Shubha Govind

## Abstract

The wasps *Leptopilina heterotoma* parasitize and ingest their *Drosophila* hosts. They produce extracellular vesicles (EVs) in the venom that are packed with proteins, some of which perform immune suppressive functions. EV interactions with blood cells of host larvae are linked to hematopoietic depletion, immune suppression, and parasite success. But how EVs disperse within the host, enter and kill hematopoietic cells are not well understood. Using an antibody marker for *L. heterotoma* EVs, we show that these parasite-derived structures are readily distributed within the hosts’ hemolymphatic system. EVs converge around the tightly clustered cells of the posterior signaling center (PSC) of the larval lymph gland, a small hematopoietic organ in *Drosophila*. The PSC serves as a source of developmental signals in naïve animals. In wasp-infected animals, the PSC directs the differentiation of lymph gland progenitors into lamellocytes. These lamellocytes are needed to encapsulate the wasp egg and block parasite development. We found that *L. heterotoma* infection disassembles the PSC and PSC cells disperse into the disintegrating lymph gland lobes. Genetically manipulated PSC-less lymph glands remain non-responsive and largely intact in the face of *L. heterotoma* infection. We also show that the larval lymph gland progenitors use the endocytic machinery to internalize EVs. Once inside, *L. heterotoma* EVs damage the Rab7- and LAMP1-positive late endocytic and phagolysosomal compartments. Rab5 maintains hematopoietic and immune quiescence as *Rab5* knockdown results in hematopoietic over-proliferation and ectopic lamellocyte differentiation. Thus, both aspects of anti-parasite immunity, i.e., (a) phagocytosis of the wasp’s immune-suppressive EVs, and (b) progenitor differentiation for wasp egg encapsulation reside in the lymph gland. These results help explain why the lymph gland is specifically and precisely targeted for destruction. The parasite’s simultaneous and multipronged approach to block cellular immunity not only eliminates blood cells, but also tactically blocks the genetic programming needed for supplementary hematopoietic differentiation necessary for host success. In addition to its known functions in hematopoiesis, our results highlight a previously unrecognized phagocytic role of the lymph gland in cellular immunity. EV-mediated virulence strategies described for *L. heterotoma* are likely to be shared by other parasitoid wasps; their understanding can improve the design and development of novel therapeutics and biopesticides as well as help protect biodiversity.

**Author summary:** Parasitoid wasps serve as biological control agents of agricultural insect pests and are worthy of study. Many parasitic wasps develop inside their hosts to emerge as free-living adults. To overcome the resistance of their hosts, parasitic wasps use varied and ingenious strategies such as mimicry, evasion, bioactive venom, virus-like particles, viruses, and extracellular vesicles (EVs). We describe the effects of a unique class of EVs containing virulence proteins and produced in the venom of wasps that parasitize fruit flies of *Drosophila* species. EVs from *Leptopilina heterotoma* are widely distributed throughout the *Drosophila* hosts’ circulatory system after infection. They enter and kill macrophages by destroying the very same subcellular machinery that facilitates their uptake. An important protein in this process, Rab5, is needed to maintain the identity of the macrophage; when Rab5 function is reduced, macrophages turn into a different cell type called lamellocytes. Activities in the EVs can eliminate lamellocytes as well. EVs also interfere with the hosts’ genetic program that promotes lamellocyte differentiation needed to block parasite development. Thus, wasps combine specific preemptive and reactive strategies to deplete their hosts of the very cells that would otherwise sequester and kill them. These findings have applied value in agricultural pest control and medical therapeutics.

## Introduction

Parasitoid (parasitic) wasps have an obligatory relationship with their insect hosts. Engaged in a biological “arms race,” each partner continuously adapts to the other to emerge alive. For reproductive success, parasitic wasps target their hosts’ behavior, development and immune system. Their attack mechanisms range from biochemical warfare and mimicry, to passive evasion and active immune suppression [1-3]. *Drosophila* and their parasitic wasps are an emerging model for studying how wasps evade or suppress host defenses [4, 5]. The generalist *Leptopilina heterotoma* (*Lh*) succeeds on the *Drosophila* species within and beyond the melanogaster group. Its close relative, *L. boulardi* (*Lb*), considered a specialist, mainly infects flies of the melanogaster group. Both wasps are highly successful on *D. melanogaster*; they consume its developing larval and pupal stages to emerge as free-living adults [6].

Oviposition into second-to-early-third *D. melanogaster* larval hosts by *Lb* and *Lh* wasps yields divergent immunological effects. *Lb* infection activates many components of humoral and cellular immunity: Toll-NF-κB, JAK-STAT, and the melanization pathways and their target genes are transcriptionally upregulated; there is a burst of hematopoietic proliferation and differentiation of blood cells (also called hemocytes) in the lymph gland and in circulation. If the immune responses are strong and sustained, macrophages and lamellocytes encapsulate and kill wasp eggs [7-11]. *Lh* infection in contrast suppresses immune gene expression and kills immature and mature larval hemocytes [12, 13].

*Lb* and *Lh* females (and also *L. victoriae*, a sister species of *Lh*) produce discrete immune-suppressive extracellular vesicle- (EV) like structures in their venom glands (called multi-strategy extracellular vesicles, MSEVs in *Lh* and venosomes in *Lb*) [14, 15]. Previously called virus like particles [16-18] these EVs lack clear viral features [15, 19]. They are produced in the venom gland, a structure made up of the long gland and a reservoir. The secretory cells of the long gland synthesize and secrete proteins some of which are initially incorporated into discrete non-spiked vesicle-like structures. In sister species *Lh* and *Lv*, these structures mature in the reservoir and assume a stellate morphology with 4-8 spikes radiating from the center. Mature EVs are roughly 300 nm in diameter, [14, 16, 20-22]. Packed with more than 150 proteins, EVs are, in part, responsible for divergent physiological outcomes in infected hosts [15, 19, 23].

Among the most abundant in the *Lh* EV proteins is a 40 kDa surface/spike protein (SSp40) [20]. SSp40 shares structural similarities with the IpaD/SipD family of proteins of the gastroenteric disease-causing Gram negative bacteria, *Shigella* and *Salmonella* [15]. Similar to SSp40’s localization to *Lh* EV spike tips, IpaD localizes to the tips of the T3 secretion injectisome, a bacterial transfer system that injects virulence proteins into mammalian cells. IpaD itself promotes apoptosis of mammalian macrophages [24, 25]. These parallels between *Lh* SSp40 and bacterial IpaD/SipD suggest that *Lh* EVs may share some similarities with bacterial secretion systems. Comparative transcriptomic/proteomic approaches revealed that SSPp40 and a few other EV proteins are not expressed in the *Lb* venom [15].

Whereas *Lh* EVs lyse lamellocytes within a few hours of wasp attack, *Lb* EVs do not have the same effect [7, 16, 26]. Our immune-inhibition experiments suggested that *Lh*’s SSp40 mediates EV interactions with lamellocytes [20]. *Lh* infection also uniquely promotes apoptosis of larval macrophages and of lymph gland hemocytes [13]. Macrophages make up more than 95% of all hematopoietic cells while differentiated lamellocytes are rarely found in naïve hosts [8-10, 27]. Work in the field strongly suggests that the protein activities concentrated within the *Lh* EVs are responsible for the destruction of these mature and immature blood cells. However, how a macro-endoparasite targets the hematopoietic system and accesses its progenitor population has not been studied. The modes of *Lh* EV entry into these cells and the pathways of destruction are also not well understood.

The goal of this study was to obtain a macro-level view of *Lh* EV interactions with cells of the larval hemolymphatic system after infection. The term hemolymph refers to the interstitial fluid that distributes hormones, peptides and other macromolecules into organs through the pumping action of an unbranched tubular heart, or the dorsal vessel. Dozens of macrophages circulate in the hemolymph. The heart lumen is surrounded by a column of paired cardiomyocytes and associated pericardial cells. This tubular structure is held in place by alary muscles [28-30]. Hematopoietic cells are organized in paired cell clusters (or lobes) on the dorsal vessel. In third instar larvae, the anterior-most lobes have blood cells at various stages of differentiation; the least differentiated progenitors are confined adjacent to the dorsal vessel, whereas the developing macrophages are sequestered in the cortical regions of the lobes (Fig 1A). In naïve hosts, the progenitor state is maintained by a putative niche (also called the posterior signaling center, PSC). The PSC is a tight unit of about 25-50 cells and is positioned posteriorly to the progenitors [31, 32]. Upon *Lb* infection, the PSC reprograms hematopoiesis inducing macrophage and lamellocyte differentiation [30, 32-39]. The entire structure is covered by the acellular basement membrane [28, 40].

**Fig 1.**
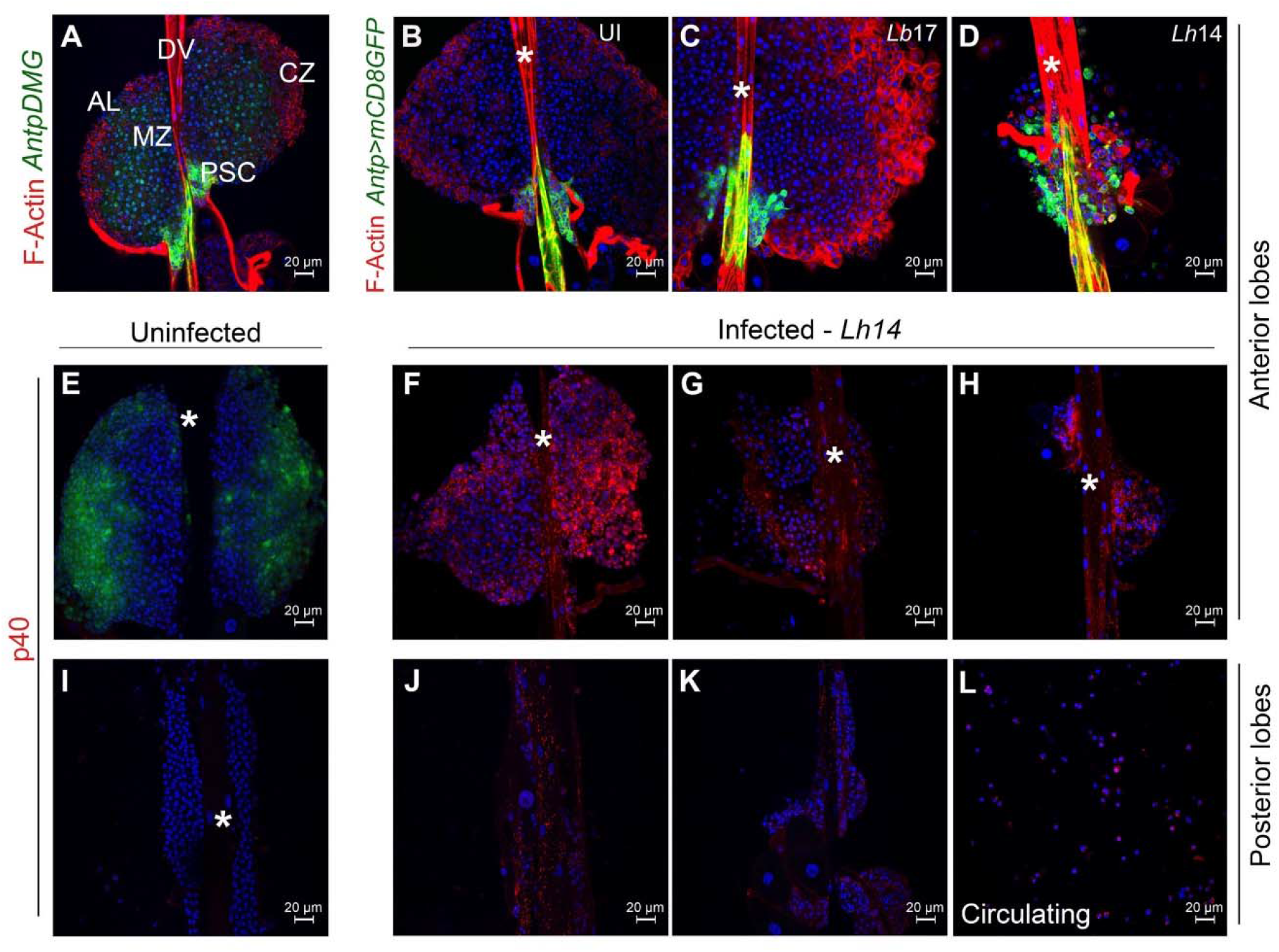
*Lh* EV association with the larval lymphatic system. (**A**) Anterior lobes (ALs, without lamellocytes) from a naive *Antp>mCD8GFP Dome-MESO-GFP* (*AntpDMG*) animal shows medullary zone (MZ), cortical zone (CZ) and posterior signaling center (PSC). The lobes flank the dorsal vessel (DV, asterisk in other panels). (**B-D**) Anterior lobes from uninfected (UI; **B**), *Lb*- and *Lh*-infected (**C, D**) *Antp>mCD8GFP* animals. *Lb17* infection induces lamellocyte differentiation in the cortex (lamellocytes are larger than their progenitors and are rich in F-actin). The GFP-positive PSC appears unaffected. *Lh14* attack leads to loss of lobe cells; the PSC cells are not tightly-clustered and displaced from their original location. (**E-K**) Lymph glands from *Pxn>GFP* animals. (**E**) GFP is expressed in the cortex of uninfected animals, but GFP expression is reduced after wasp attack (**F-H**). *Lh14*-infected ALs from the same infection experiment showing variability in loss of cells; *Lh* EVs (anti-SSp40 staining, referred to as p40 here and remaining figures) are seen in some cells of these lobes and within the dorsal vessel. (**I-K**) Posterior lobes from uninfected (**I**) and *Lh*-infected animals (**J, K**). (**L**) Circulating hemocytes from infected animals.

Using *Drosophila* genetics, cell-specific markers and SSp40 staining as a proxy for *Lh* EV localization, we have pieced together a broad view of this host-parasite interaction interface. We show an abundance of *Lh* EVs in (a) the lumen of the larval dorsal vessel, (b) along the collagen/perlecan-based basement membrane around the dorsal vessel and surrounding clusters of lymph gland progenitors, and (c) inside the progenitor and mature macrophages. Moreover, high EV signal correlates with the disassembly of the cohesive PSC unit. PSC ablation limits EV internalization and loss of lymph gland hemocytes, while PSC inactivation via *hedgehog* (*hh*) knockdown (KD) does not have this effect. We also show that lymph gland hemocytes can phagocytose *Lh* EVs using the classical Rab5-mediated retrograde transport (RGT) pathway. Surprisingly, Rab5 also maintains fly macrophage identity, as *Rab5* knockdown leads to over-proliferation, lamellocyte differentiation and tumorigenesis; *Rab5* function is cell-autonomous. Thus lymph glands are not merely a source of mature blood cells but are themselves immune competent organs and can clear the immune-suppressive *Lh* EVs to defend the host. However, *Lh* EVs proactively dislodge cells of the PSC, blocking differentiation of the protective immune cells. *Lh* EVs target the endomembrane system of macrophages that ultimately results in their apoptosis, thus highlighting central and previously unrecognized roles of the lymph gland in cellular immunity. These observations help explain why *Lh* infections target the larval lymph gland. The direct EV-macrophage interactions and cellular outcomes set the stage for future molecular analyses in both the hosts and parasites.

## Results

### Lh EVs are present within the larval lymphatic system

*Lb17* attack triggers lamellocyte differentiation in the larval lymph gland cortex (Fig. 1A-C). At an equivalent time-point, *Lh*-infected lobes are significantly smaller (Fig. 1D-K); [13]. Surprisingly, unlike *Antp>mCD8GFP*-expressing PSCs of naive and *Lb-*infected lobes that remain tightly clustered (Fig. 1A-C), PSC cells of *Lh*-infected hosts are dislodged and some are distributed in the body of the lobe (Fig. 1D).

To understand these responses, we imaged more than 25 hosts in multiple experiments. Throughout these studies, we used a polyclonal antibody to mark SSp40, an *Lh* EV-specific protein [20]. *Lh* infection of *Pxn>GFP* animals reduced *Pxn>GFP* expression (Fig. 1E-H). (*Pxn* is normally active in the cortex and its expression mimics that of many other genes downregulated by *Lh* infection [7]). An abundance of *Lh* EVs was observed in anterior- and posterior-lobe hemocytes as well as in the dorsal vessel (Fig. 1F-H, J, K). This staining signal is absent in glands from naive animals (Fig. 1E, I). Thus, *Lb* and *Lh* attack have drastically different outcomes and *Lh* EVs appear to interact directly with most lymph gland hemocytes.

In our analyses across multiple experiments, we found that the degree of tissue loss and EV distribution varies. Lobe morphologies ranged from nearly intact and filled with EVs (Fig. 1F, J) to damaged lobes and many or few EVs (Fig. 1G, H, K). This variation is likely due to (a) the duration of infection (i.e., time between oviposition and dissection); (b) the injected EV dose; or (c) the dynamics of EV circulation. Dissections at later time points showed loss of almost all lymph gland hemocytes [13].

To then probe how *Lh* EVs enter the lymphatic system, we stained *Lh*-infected glands from fly strains with GFP-tagged Collagen IV (basement membrane, *Viking* [41]) or GFP-tagged proteoglycan core protein, Perlecan/Trol [42]. In both cases, EV puncta were clearly localized with the continuous GFP signals of these extracellular matrix (ECM) proteins along the dorsal vessel as well as in the interstitial spaces around clustered hematopoietic progenitors (yellow puncta in Fig. 2A-D’ arrows). Surprisingly, punctate staining was also seen inside immature progenitors, adjacent to the dorsal vessel (Fig. 2B, B’ D, D’). EVs were also observed inside some cardiomyocytes as evidenced by SSp40 colocalization with the mCD8GFP signal in *HandΔ>mCD8GFP* larvae (Fig. 2E-F’, arrows). Thus, *Lh* EVs associate with ECM proteins of the lymphatic system, enter the dorsal vessel lumen and even some cardiomyocytes.

**Fig 2.**
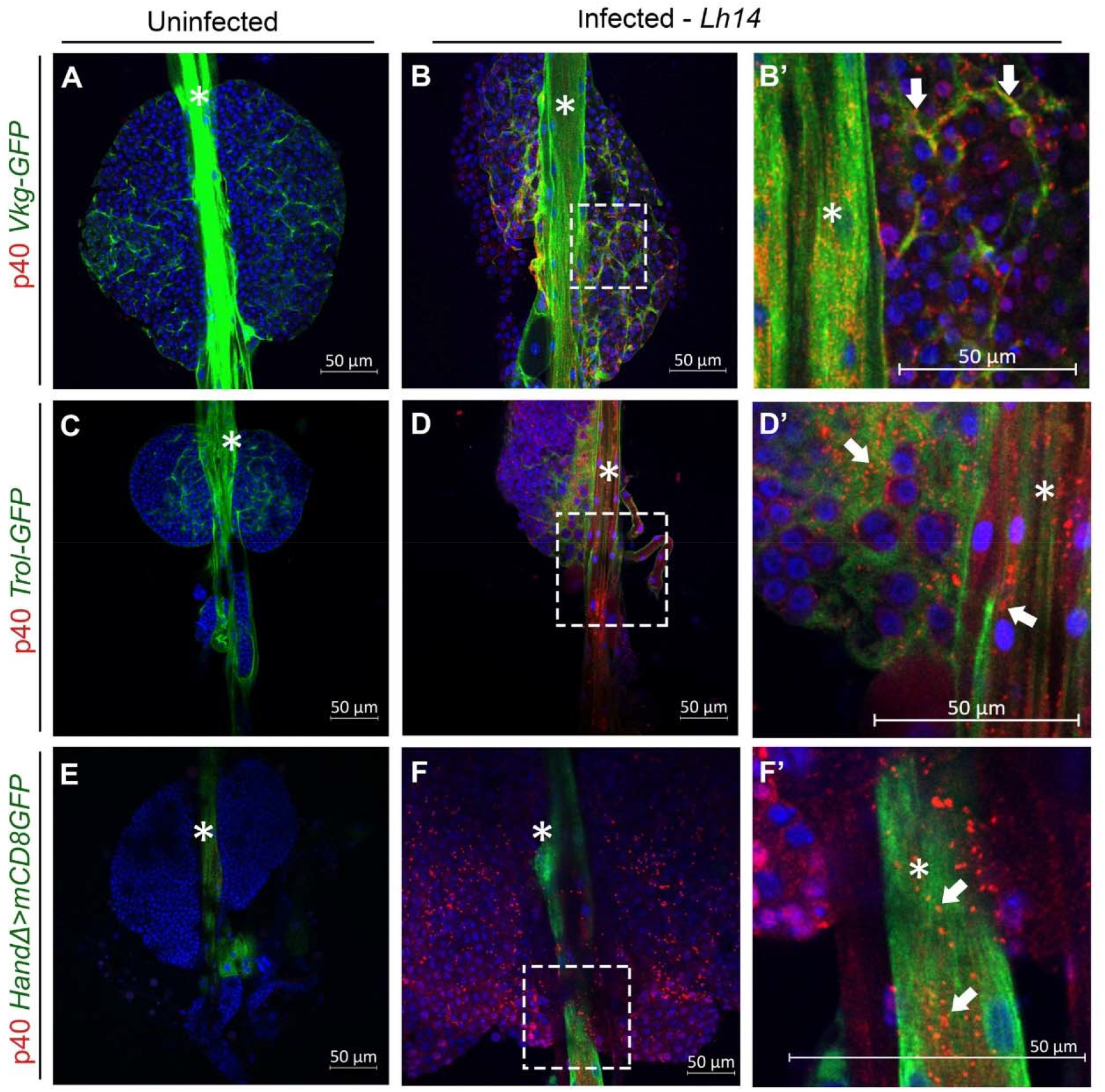
*Lh* EVs association with basement membrane proteins. (**A-B**) *Vkg-GFP* lymph glands. (**A**) GFP marks Collagen IV in the dorsal vessel and around lobe clusters. (**B-B’**) *Lh* infected gland shows extensive EV puncta co-localized with GFP in the dorsal vessel (*) and in between lobe clusters. Punctate, cytoplasmic staining in many cells throughout the anterior lobes (B, B’, arrows) is also observed. (**C-D**) *Trol-GFP* lymph glands. (**C**) GFP marks Trol-GFP distribution in naïve and (**D, D’**, arrows) *Lh* infected animals. Co-localization of *Trol-GFP* and EV puncta are observed. (**E-F**) *HandΔ>mCD8GFP* animals marks the cells of the dorsal vessel. (**E**) An intact lymph gland from a naïve animal. (**F, F’**, arrows) EV signal localizes with the GFP signal in the cardiomyocytes.

### Effects of Lh infection on PSC integrity

Previous studies have demonstrated that upon *Lb* infection, the PSC reprograms hematopoietic development and promotes lamellocyte differentiation. Similar to controls, *Lb* infected PSCs remain tightly clustered [35, 37, 38, 43, 44]. We found that regardless of the *Lh* strain, PSC cells dislodged from their normal posterior position into the body of the lobe (Fig. 1D and Fig. S1A, B). At the time points we examined, half of the PSC cells relocated into the lobe, of which an overwhelming majority (>95%) were present as single cells or as groups of two cells. The other half remained in their original place, although some were not as tightly clustered (Fig. 1D; Fig. S1; n=24 PSCs from 12 lymph glands). As expected, in 14 lobes from naive controls, all PSC cells were tightly packed. Strikingly, in samples where the PSCs were still intact, EVs congregated in regions adjacent to the PSC, but were never found inside the PSC cells (Fig. S1A, B).

The Slit ligand, originating from adjacent cardiomyocytes, controls PSC integrity via the Robo receptors in the PSC; Robo2 has the strongest effect [45]. Indeed, the effect of *Lh* infection on the PSC resembles Slit/Robo2 knockdown, which promotes PSC disassembly [45] (Fig. 1 and Fig. S1). A cohesive PSC is important in hematopoietic development as fewer macrophages and crystal cells develop in *Slit*/*Robo2* KD lobes compared to controls. But differing from *Lh* infection, *Slit* KD PSCs are larger and there is no apparent loss of progenitors [45]. In spite of different outcomes in the two conditions, we hypothesized that the initial steps might be shared and that *Lh* EVs might inactivate the Slit-Robo signal, which might explain PSC disassembly.

To test this idea, we infected animals in which the Slit-Robo pathway was manipulated to promote constitutive signaling. We found that neither expressing active Slit nor overexpressing Robo2 altered *Lh* EVs’ ability to disassemble the PSC. *Lh* infection of *HandΔ>Slit-N* animals still promoted PSC disassembly (Fig. S2A-C’; n =16 lobes) even though gain-of-function *Slit-N* [46] reverses the effects of Slit KD [45]. Similarly, *Lh* infection bypassed the effects of Robo2 overexpression (*Antp>Robo2-HA*) and promoted PSC disassembly Fig. S2D-F’). Many EVs are observed around these PSCs (Fig. S3; n=16). Thus, either EVs inactivate PSC function independently of the Slit-Robo signal, or they possess redundant mechanisms that disable constitutive Slit-Robo signaling.

### PSC-less lymph glands remain intact

PSC-less lymph glands were unable to induce lamellocyte differentiation after *Lb* infection [36]. We asked if ablating the PSC might similarly inhibit the *Lh* infection responses. PSC-less lobes (*Col>Hid*) lacked Antp staining and *Lh14* infection did not affect lobe integrity (Fig. S4A-D’; n>12 lobes). Moreover, while *Lh*-infected *UAS-Hid* lobes lost progenitors and exhibited high levels of EV uptake (Fig. 3A, B, D, E), *Lh*-infected PSC-less lobes remained intact and showed low, a non-specific SSp40 staining signal in anterior and posterior lobes (Fig. 3C, F; n>12 lobes for each condition). In contrast to the non-responsive *Col>Hid* lobes, *Antp>hh*^*RNAi*^ lobes responded to *Lh* infection and suffered progenitor cell loss (Fig. S5A-F; n>12 lobes for each condition). Thus, inactivating the signaling function of the PSC does not appear to affect the wasp’s ability to disassemble the PSC. P1 staining revealed that *Lh* infection does not block macrophage differentiation (Fig. S5C). Taken together, these results suggest that the PSC plays a structural role in trafficking of EVs from either the hemolymph or the dorsal vessel into the lobes and that the cell-lethal effects of *Lh* EVs is distinct or downstream of the PSC’s niche function.

**Fig 3.**
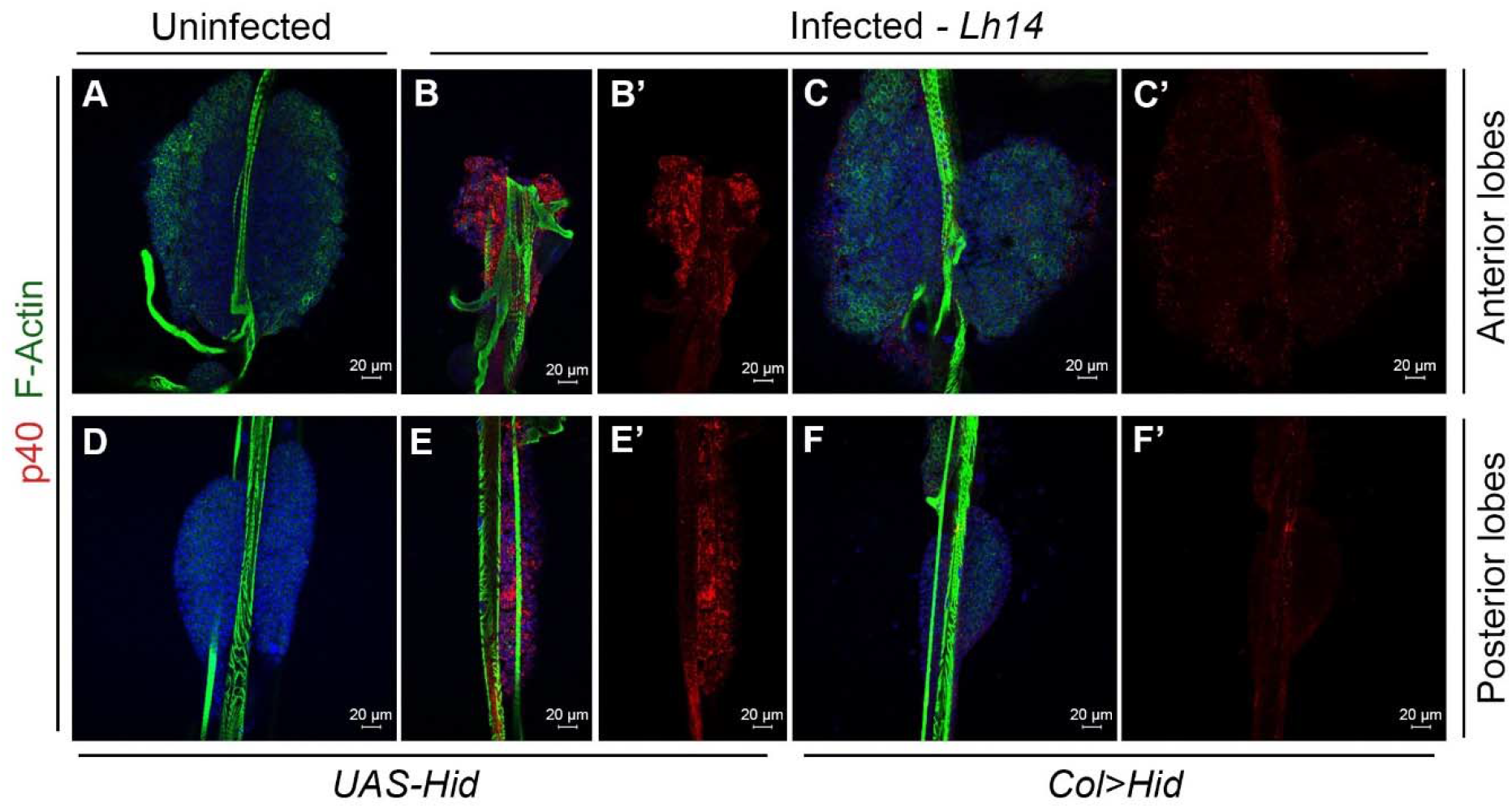
PSC-less lymph glands from *Lh-*infected animals have intact lobes and show reduced EV signal. (**A, D**) Anterior lobes from a naïve *UAS-Hid* host are intact and do not have EVs; anterior (**A**), and posterior lobes (**D**). (**B, E**) Anterior lobes from a *Lh* infected *UAS-Hid* host is depleted of hemocytes and has many EVs. Anterior lobes (**B, B’**), and posterior lobes (**E, E’**). (**C, F**) PSC-less lobes from a *Lh* infected *Col>Hid* host are intact with weak EV signal. Anterior (**C, C’**), and posterior lobes (**F, F’**).

### The larval lymph gland is phagocytically competent

We next studied if EV uptake into hematopoietic cells occurs via RGT mechanisms. In *Pxn>GFP*; *+/Bc* heterozygous lymph glands, we observed blackened, dead crystal cells in the cytoplasm of the GFP-positive cortical cells (Fig. S6). This observation suggests that GFP-positive lymph gland cells are phagocytically competent. We therefore investigated if *Lh* EV uptake depends on Rab5, an early endosomal protein. Rab5 mediates trafficking from the plasma membrane to early endosomes [47]. In contrast to *Pxn>GFP* macrophages, where EV staining is bright and punctate throughout the cytoplasm (Fig. 4A, B), *Pxn>GFP, Rab5*^*RNAi*^ cells show peripheral punctate staining, presumably from intact EVs, trapped in early endosomes, both in lymph gland (Fig. 4C; arrows) and circulating (Fig. 4D; arrows) hemocytes. In lamellocytes, the EV signal is diffuse and nuclear, and Rab5 KD shows no change in staining intensity or distribution (Fig. 4E, F; arrowheads), suggesting Rab5-independent uptake mechanisms are involved. *msn-GFP*-and Integrin-β-positive lamellocyte fragments were also observed in *Lh*-infected macrophages suggesting occurrence of efferocytosis (Fig. S7).

**Fig 4.**
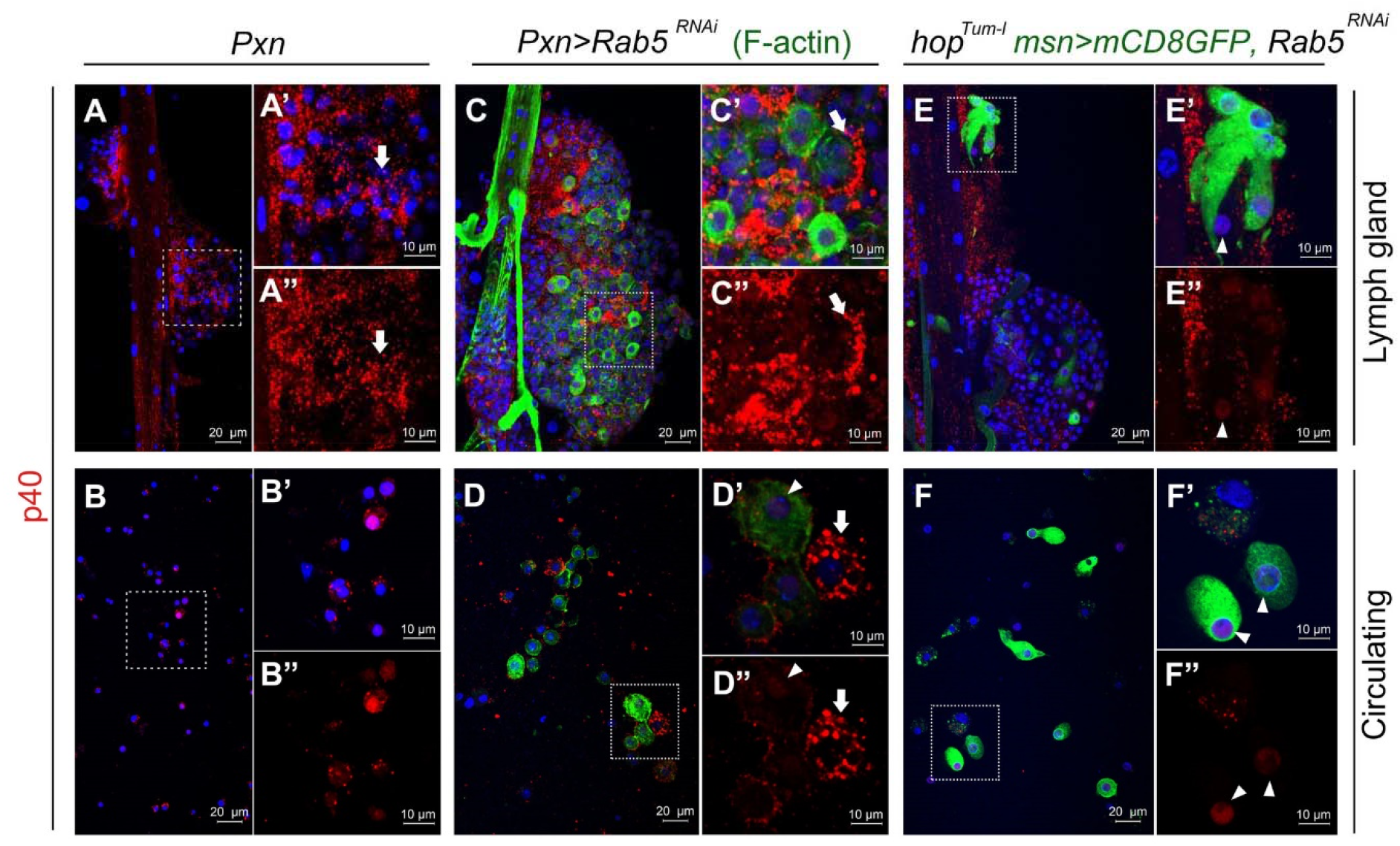
Intracellular *Lh* EV localization. (**A, B**) Anterior lobes (A-A”) and circulating hemocytes (B-B”) from *Lh*-infected *Pxn>GFP* animals showing EV uptake. Magnifications of areas in (A) and (B) are shown in A’, A” and B’, B”, respectively. Arrows point to internalized vesicles. (**C, D**) An anterior lobe (C-C”) and circulating hemocytes (D-D”) from *Lh*-infected *Pxn>GFP Rab5*^*RNAi*^ animals showing peripheral localization of EVs (arrow). As in Fig. 1, *Pxn>GFP* expression is reduced after wasp attack. Samples were counterstained with FITC-Phalloidin to visualize cell morphology. A larger Phalloidin-positive lamellocyte (arrowhead) remains EV negative, while smaller macrophages endocytose EVs. The EV signal is peripheral in some, but not all cells. (**E, F**) Lobes (E-E”) and circulating hemocytes (F-F”) from *Lh*-infected GFP-positive lamellocytes of *hop*^*Tum-l*^ *msn>mCD8GFP, Rab5*^*RNAi*^ animals. Lamellocytes show a diffuse nuclear SSp40 signal (arrowhead).

### Lh EVs negatively impact phagolysosomal organization in macrophages

Rab7 mediates late endosome formation and trafficking between late endosomes and lysosomes, marked by Rab7 and LAMP1, respectively [47]. To evaluate if *Lh* EVs impact the RGT machinery, infected glands expressing GFP-tagged Rab5, Rab7, or LAMP1 proteins were examined (Fig. 5). Under our experimental conditions, *Lh* EVs rarely colocalized with early endosomes and Rab5 compartment morphology remained comparable to uninfected controls (only 14% of SSp40 puncta are Rab5-positive; n = 221 cells; 6 lobes; Fig. 5A-C). In contrast, *Lh* EVs were consistently found with GFP-Rab7 and GFP-LAMP1 and these compartments were grossly distorted (100% co-localization; n = 115 and 112, Rab7 and LAMP, respectively, 6 lobes each; Fig. 5D-E”). Moreover, the Rab7/EV and LAMP1/EV signals were asymmetrically localized in *Lh*-affected cells. These observations suggest that high numbers of *Lh* EVs transit through early endosomes, but that they are retained in late RGT compartments including lysosomes. Thus, *Lh* EVs have a detrimental effect on RGT compartment integrity and this loss of integrity may promote lysosomal leakage and labilization.

**Fig 5.**
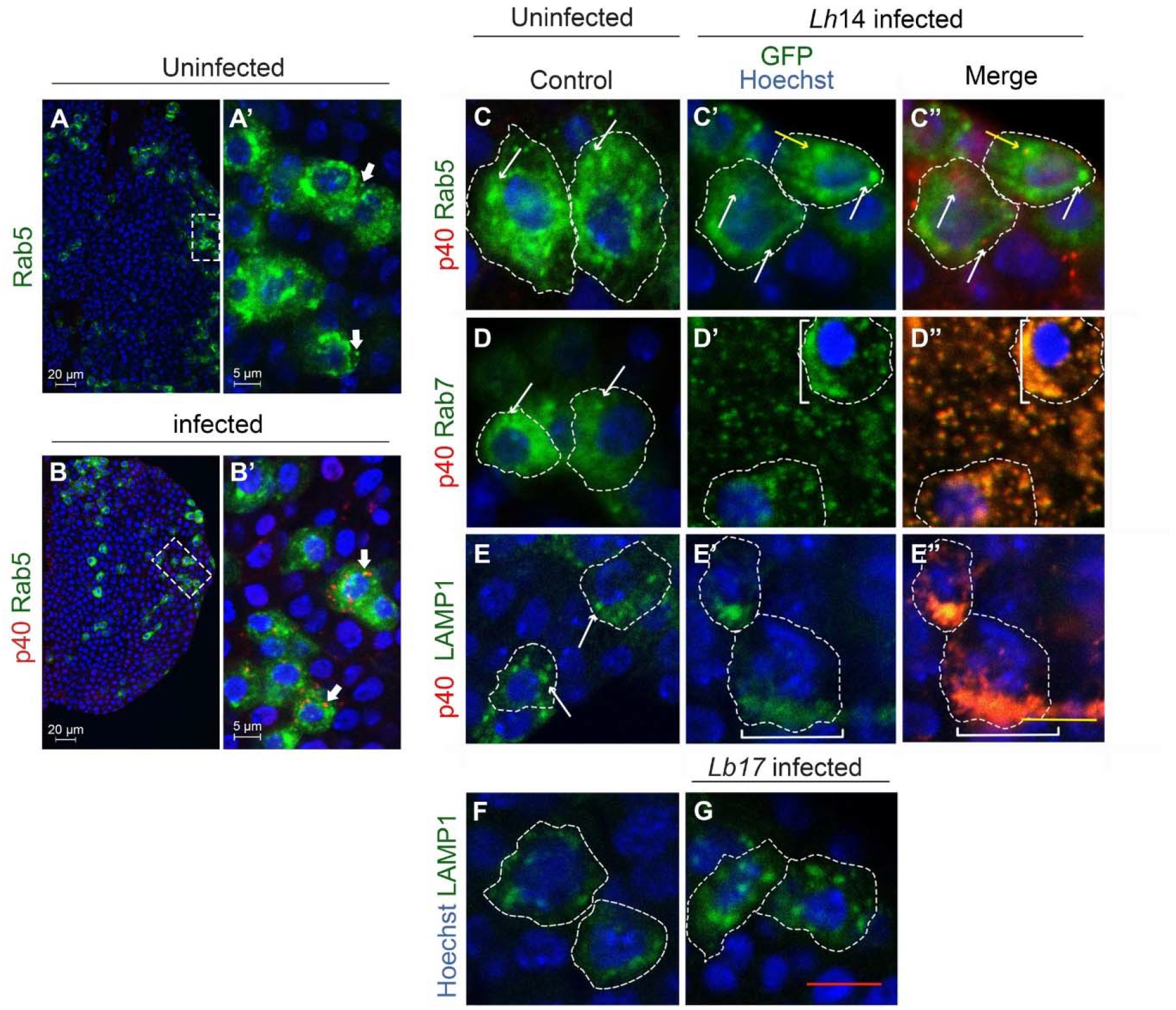
Effects of *Lh* and *Lb* infection on retrograde transport organelles. (**A, B**) *Hemese>GFP-Rab5* lymph gland lobes stained with anti-SSp40; *Lh* EVs enter Rab5 compartments, some EVs colocalize with GFP-Rab5 (arrows). (**C-E**) *Hemese>GFP-Rab5, >GFP-Rab7*, and *>GFP-LAMP1* expression in naïve animals or (C – E prime) *Lh*-infected animals. Individual cells are outlined. Yellow arrow in C’ and C” shows a normal Rab5 compartment with EV signal. White arrows point to normal compartment morphologies. Many *Lh* EVs are associated with grossly distorted Rab7 and LAMP1 compartments (square brackets). (**F, G**) *Lb* infection does not distort LAMP1 compartment morphologies.

### Rab5 suppresses proliferation and maintains the macrophage fate

We were surprised to find that *Pxn>GFP Rab5*^*RNAi*^ animals developed melanized tumors (Fig. S8A, B); the hematopoietic population is significantly expanded and lamellocyte differentiation is robust, affecting viability (Fig. S8C-F). (Variability in viability and tumor development in Rab5 KD animals is likely due to the strength and differential cell-specific expression patterns of the GAL4 drivers.) A similar result was observed with the expression of the dominant negative Rab5S34N protein which cannot bind GTP [48]). Rab5 KD even in the lymph gland medullary zone (*TepIV>Rab5*^*RNAi*^) resulted in lamellocyte differentiation (Fig. 6A, B).

**Fig 6.**
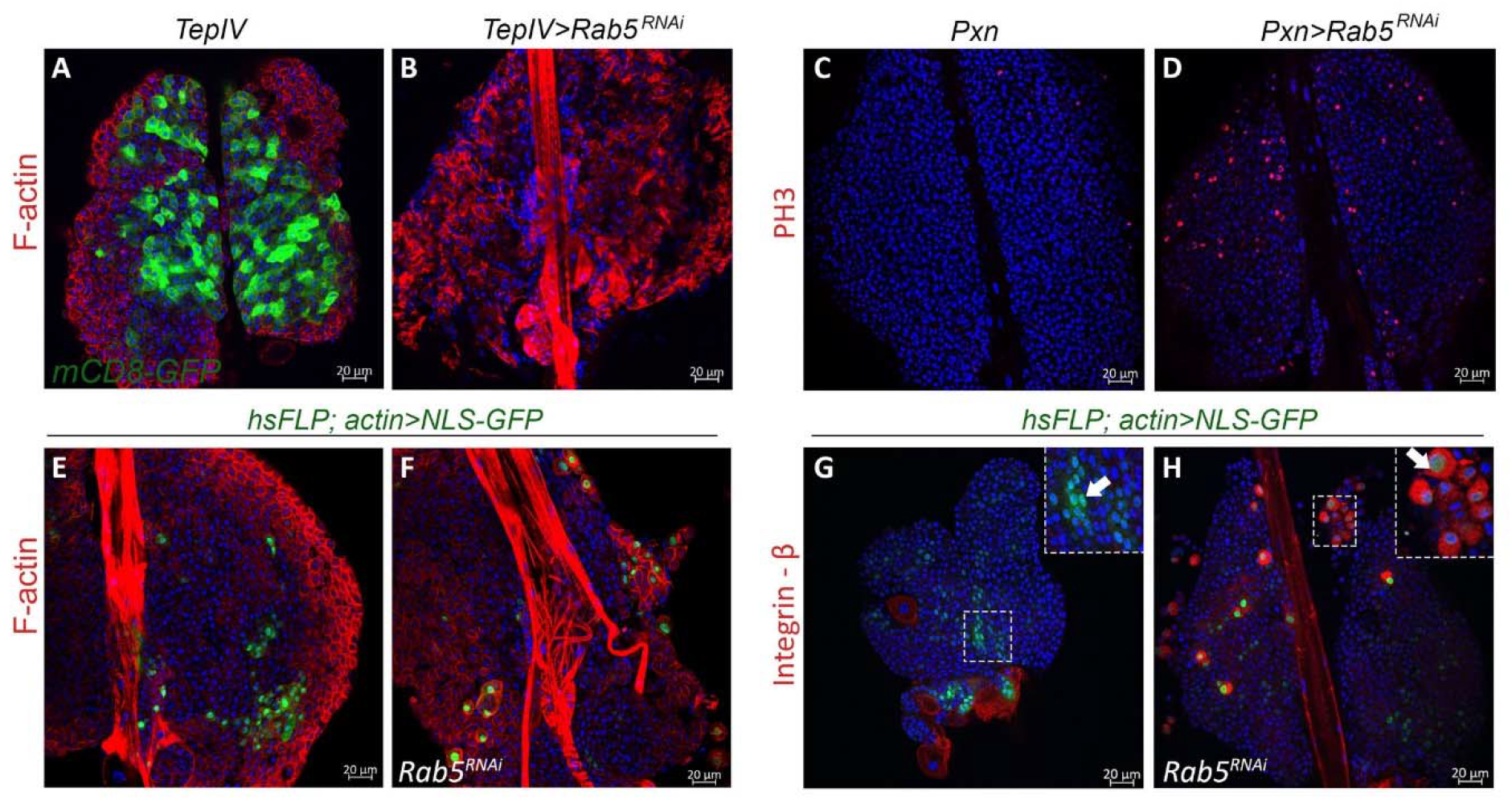
*Rab5*^*RNAi*^ triggers overproliferation and lamellocyte differentiation. (**A, B**) *TepIV>Rab5*^*RNAi*^ in the medullary zone drives lamellocyte differentiation. Lamellocytes are rich in F-actin. (**C, D**) High levels of mitosis (PH3-positive) are induced by *Pxn>Rab5*^*RNAi*^. (**E-H**) Cell-autonomous inhibitory role for *Rab5* in lamellocyte differentiation. (**E, G**) *hsFLP; actin>NLS-GFP* control clones (without Rab5 KD) are marked with GFP and do not contain lamellocytes (inset, arrow in panel **G**). (**F, H**) *hsFLP; actin>NLS-GFP, Rab5*^*RNAi*^ clones are composed of lamellocytes with high F-actin staining (**F**) and are Integrin-β-positive (**H**). In panel H, the GFP and Integrin-β signals overlap in some cells confirming lamellocyte identity (inset, arrow in panel **H**).

Hematopoietic expansion correlated with increased mitotic index (MI) in lobes of tumor-bearing *Pxn>GFP Rab5*^*RNAi*^ animals (Fig. 6C, D) suggesting that normal Rab5 function checks over-proliferation and ectopic progenitor differentiation. (MI=2.2 ± 2.15 in control *Pxn>GFP*; 2.9 ± 1.1, and 5.5 ± 1.9 in experimental *Pxn>GFP, Rab5*^*RNAi*^ animals without and with tumors, respectively (n = 10 for each condition). These results suggest that *Rab5* acts as a tumor suppressor and maintains hematopoietic immune quiescence.

Control “FLP-out” clones without *Rab5*^*RNAi*^ contained small cells that did not express integrin-β; experimental clones with *Rab5*^*RNAi*^ had larger, F-actin-rich cells with a typical lamellocyte morphology, that were also integrin-β-positive (Fig. 6E-H). These results suggest that Rab5’s requirement in maintaining the progenitor or macrophage fate is cell-autonomous.

## Discussion

### System-wide distribution but specific effects of *Lh* EVs

Parasitism by *L. heterotoma* has been of interest because of its ability to parasitize many *Drosophila* hosts and the existence of venom factors that kill host hemocytes. The discovery of an anti-lamellocyte activity intrinsic to *Lh* EVs provided initial insights into the critical roles of these EVs in parasitism [20, 49]. However, details underlying their apoptotic effects on macrophages have been lacking. This work provides the first view into how *Lh* EVs rely on the host’s circulation to gain system-wide distribution to not only precisely kill the available effector cells but also to pre-emptively interfere with the host’s ability to produce additional effector cells. We show that lymph glands serve an important, previously unappreciated role in immunity. A majority of lymph gland cells can phagocytose *Lh* EVs to protect the host from their detrimental effects. EV activities in turn promote their apoptotic death by disrupting their endomembrane system.

This work also lays bare new questions. *Lh* EVs’ association with the ECM proteins around the lymphatic system cells suggests ways in which EVs might recognize and home into the lymph gland hemocytes and cardiomyocytes although the role of the ECM, the details of their entry and physiological effects on cardiomyocytes are currently unclear. As has been suggested for Slit-carrying vesicles [45], cardiac cells might provide a route for *Lh* EVs to converge into the vicinity of the PSCs. Once inside lymph gland hemoctytes, they simultaneously target the protective functions of macrophages and lamellocytes and their activities culminate to strongly block encapsulation (Fig. 7). These strategies are not uncommon and are likely to be shared by closely related *Leptopilina* wasps or even unrelated virulent wasps that attack drosophilid and non-drosophilid hosts and are known to destroy their hosts’ hematopoietic cells [50-54].

**Fig 7.**
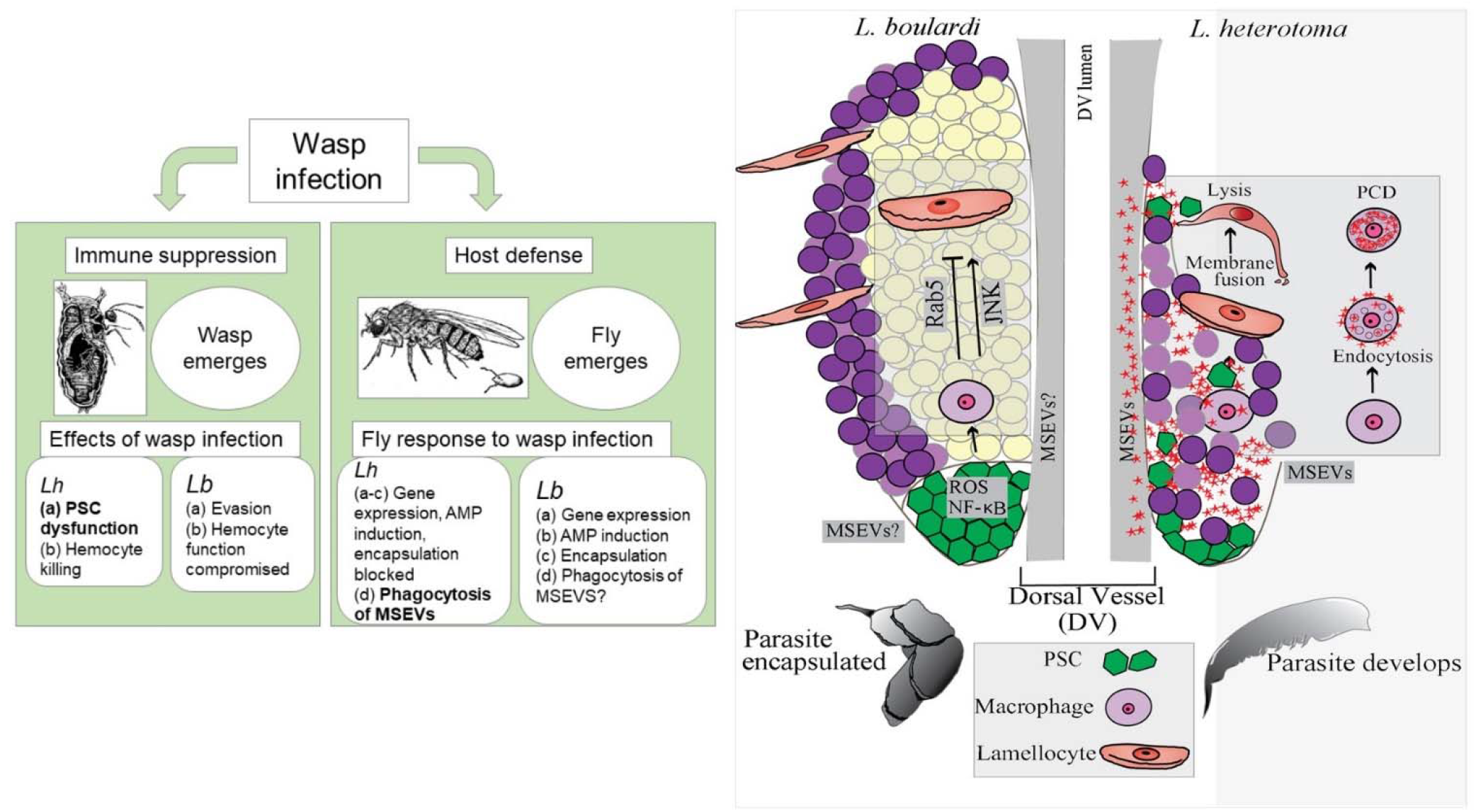
*Lh* EV interactions and effects on host blood cells: summary of events. Left: Immune suppression by *Lh* and *Lb* wasps and host defense in response to infection. Text in bold indicates findings from this study. Right: *Lb* (left lobe) attack triggers signaling events in the PSC and promotes lamellocyte differentiation of lymph gland progenitors. This process is important in host defense against parasitic wasps and is kept in check by Rab5. After *Lh* infection (right lobe), *Lh* MSEVs concentrate around and disassemble the PSC. They are phagocytosed by macrophages by a Rab5-dependent endocytic mechanism. In macrophages, EVs damage the phagolysosomal compartments. They are internalized by lamellocytes independently of Rab5 function. EVs lyse the few lamellocytes that differentiate post infection.

### *Lh* EVs proactively block encapsulation

Studies with *Lb* showed that the lymph gland itself responds to wasp infection and lamellocytes differentiate from hematopoietic progenitors [8, 9, 55, 56]. In addition to PSC’s niche function in naïve animals [31, 32, 57-59], the PSC also appears to play an anti-parasite role as *Lb* infection promotes lamellocyte differentiation [30, 35-38, 43, 60]. Given this latter role, it is reasonable to interpret *Lh’*s effects on PSC integrity as part of a corresponding adaptive strategy that *Lh* has acquired during its evolutionary history.

The high *Lh* EVs levels around the PSC and its disassembly provide novel physiological insights into PSC functions and raise intriguing mechanistic questions. A normally clustered and cohesive PSC organization is needed for proper hematopoietic differentiation in naïve animals [45]. Although *Lh* infection-induced PSC disassembly and hemocyte loss are observed together in fixed samples, our data from infected PSC-less animals suggest that PSC disruption might precede hemocyte death. It is possible that the PSC may somehow “recognize” foreign entities and might serve to protect the progenitor microenvironment by acting as a chemical or mechanical barrier between the vascular and hematopoietic cells. In this scenario, *Lh* EVs may be targeting PSC cohesiveness to inactivate this barrier function.

This interpretation is consistent with the recently discovered permeability barrier in the PSC that is breached by systemic bacterial infection. Permeability barrier in the PSC is maintained by septate junctions (SJ) and SJ depletion is linked to increased Toll signaling, cellular immune activation and improved host survival [39]. The barrier function between the vascular and hematopoietic cells proposed here would serve to limit the ingress of structures such as microbes and EVs. The mechanisms underlying the effects of ablated PSCs are unclear but it is notable that PSC-less lobes do not respond to either *Lb* or *Lh* infections (our results and [61]). Examining whether *Lb* EVs/venosomes similarly interact with the lymph gland ECM, congregate around the PSC, and are phagocytosed by hemocytes will shed more light on these processes.

Extracellular vesicles secreted from mammalian neutrophils and endothelial cells can direct cell motility and chemotaxis [62, 63]. Thus it is possible that one or more *Lh* EV activities [15] is similarly responsible for PSC disassembly. These activities may perturb host pathways required for normal PSC cohesion and function, although manipulating the Slit-Robo pathway components, or *hh* signaling, hypothesized to disrupt the infection process, was insufficient to block *Lh*’s ability to disperse the PSC and attack hemocytes. *Lh* EVs may possess redundant or independent mechanisms to control PSC integrity and this question remains open for further research.

### A central role for phagocytosis in the anti-parasite response

The ability of macrophages to ingest and kill microbes is a fundamental facet of innate immunity. Microbes have evolved to evade or escape the destructive conditions in their host cells’ phagolysosomes. While most intracellular pathogens avoid fusion with lysosomes, others modify endocytic trafficking differently to survive in their host cells [64]. We have shown that like microbes, *Lh* EVs are endocytosed and can damage the late endocytic compartments. This suggests that their biochemical activities may distort and damage intracellular membranes although how this occurs is unclear. A novel family of *Lh* EV-associated GTPases [15] are possible candidates for such activities as expression of select GTPases in yeast alter vacuolar morphologies [65]. Microbial infection of macrophages can activate apoptosis responses [66-68] and it is possible that similar effects of *Lh* EVs in fly macrophages are directly linked to their apoptosis.

Lamellocytes utilize a flotillin/lipid raft dependent mechanism to internalize *Lb* EVs [69], and it is possible that *Lh* EVs use the same or a similar pathway. The significance of the nuclear SSp40 signal in lamellocytes after *Lh* infection is unclear; but because the signal is not punctate, *Lh* EVs are likely internalized via a membrane fusion step in which their vesicular character is lost. EM results also show that unlike macrophages with membrane-enclosed endocytic vesicles containing intact *Lh* EVs, lamellocytes do not have such compartments, and once internalized, EVs lose structural integrity [49]. Efferocytosis thus appears to be an effector anti-parasite response as lysed lamellocytes are cleared by this process, and it may ultimately also be beneficial to parasite development.

Our genetic studies with Rab5 highlight the central role of the endocytic processes in the anti-parasite response. Loss of endocytic trafficking activates immune signaling ([70-72]). From a physiological standpoint, it makes sense why the lymph gland is a dedicated target of wasp infections. Both key aspects of anti-wasp cellular immunity, i.e., phagocytosis of the wasp’s EVs and lamellocyte differentiation, unequivocally reside in the lymph gland. These ideas can be further examined at the molecular level with the available descriptions of the *Lh* and *Lb* EV proteomes [15, 19, 23]. Virulence factors provide the armament for parasite success in the host/pathogen arms race. Insights from this model host-parasite system can influence our understanding of how parasite-derived factors have shaped the immune physiology of fly hosts.

## Author contributions

All authors conceived the experiments, JR, MEH and ZR performed experiments; SG supervised the project and wrote the manuscript with input from coauthors.

## Competing interests

The authors declare no competing interests.

## Acknowledgements

We are grateful to the Bloomington Drosophila Stock Center, Developmental Studies Hybridoma Bank, and colleagues for provision of fly stocks and antibodies. Funding for this work came from NASA (NNX15AB42G), National Science Foundation (1121817 and 2022235), National Institutes of Health (1F31GM111052-01A1 to MEH and G12MD007603-30 to CCNY) and Howard and Vicki Palefsky Fellowship to JR. The sponsors or funders did not play any role in the study design, data collection and analysis, decision to publish, or preparation of the manuscript.

## Materials and Methods

### Stocks and crosses

All *D. melanogaster* stocks were raised on standard fly medium containing cornmeal flour, sucrose, yeast, and agar at 25°C.

#### GAL4 lines

PSC drivers were: *Antp-GAL4; mCD8GFP* ([73], from S. Minakhina) and *y w; Collier-Gal4/CyO y*^*+*^ ([32] from M. Crozatier). The truncated *HandΔ* promoter is active in cardiomyocytes of the dorsal vessel ([74], from M. Crozatier). Hemocyte drivers were: *Pxn-GAL4, UAS-GFP* ([75], from U. Banerjee); *Hemese-GAL4* (*He-GAL4*) ([76], from D. Hultmark); *eater (ea)-GAL4 (*[77], from R.A. Schulz); *Collagen-GAL4* (*Cg>GFP*) ([78], from C. Dearolf); *Serpent (Srp)-GAL4* [79] and *TepIV-GAL4* [80] (both from N. Fossett); *Hemolectin-GAL4* (*Hml>GFP*) ([81], from J-M. Reichhart).

#### UAS lines

*UAS-Slit-N* ([46]; Slit gain-of-function) and *UAS-Robo2-HA* (for overexpression of Robo2, [82]) lines were obtained from T. Volk and T. Kidd.

Strains from the Bloomington Drosophila Stock Center: *UAS-Rab5*^*RNAi*^ (#30518); *UAS-GFP-Rab5* (#43336) [83]; *UAS-GFP-Rab7* (#42706); *UAS-Rab5*.*S43N* (#42704); *UAS-GFP-LAMP*; *nSyb-GAL4/CyO:TM6B* (#42714) [84], and *UAS-hh*^*RNAi*^ (#25794) [85]).

#### Other lines

A homozygous *Bc* stock devoid of markers and other mutations (from B. Lemaitre [86]) was balanced with the *CyO-GFP* balancer for crosses with the homozygous *Pxn-GAL4, UAS-GFP* strain. Protein trap lines were: *Collagen IV* (Viking) and *perlecan* (Trol) (from A. Spradling and L. Cooley); *hhf4f-GFP; Antp-GAL4/TM6 Tb Hu* (marks PSC, from R.A. Schulz); *Dome-MESO-GFP* (lymph gland medulla is GFP-positive) ([87] from M. Crozatier). We recombined this latter insertion with the *Antp-GAL4* insertion to make a *UAS-mCD8GFP; Antp-GAL4, Dome-MESO-GFP* (*AntpDMG*) stock. *hop*^*Tum-l*^, *msn-GAL4; UAS-mCD8GFP* [88] uses the *misshapen* (*msn*) driver to mark lamellocytes [89]. For PSC-less animals, *UAS-Hid* [90] females were crossed with *Collier-GAL4/CyO y+* males. For FLP-out clones [91], *hsp70-flp; Actin>CD2>GAL4* flies were crossed with the *Rab5*^*RNA*i^ flies; progeny was heat shocked at 37°C as described [38]. UAS-GAL4 crosses were maintained at 27^°^C.

### Wasp infections

*y w* flies were used to rear wasps. Unless specified otherwise, infections were done with either *Lb17* or *Lh14* [7]. *LhNY* [20] was used to validate results with the *Lh14* strain. Ten to twelve trained female wasps were introduced to hosts from a 12-hr egg-lay. Hosts were allowed to recover after an 8-12 hr infection. Dissections were typically done two days after infection. Uninfected controls followed the same timeline. In general, longer infection regimes led to stronger responses: more lamellocytes differentiated after *Lb* infection and more lobe cells were lost after *Lh* infection. Under our experimental conditions, superparasitism by either wasp was rare and for our analyses, we avoided hosts with more than one parasite.

### Immunohistochemistry

Antibody staining was performed according to [92]. Primary mouse anti-SSp40 (1:1000) [20] and Cy3 AffiniPure donkey anti-mouse secondary (1:200) (Jackson Immuno Research) were used to detect *Lh* EVs. Mouse anti-Antennapedia (1:10; Developmental Studies Hybridoma Bank 8C11, [93] and macrophage-specific mouse anti-P1 (1:20; I. Ando [94]) were similarly detected. Nuclear dye (Hoechst 33258, Invitrogen, 1:500) and Rhodamine or Alexa Fluor 488-tagged Phalloidin (Invitrogen) were used for counterstaining cells. For mitotic index, rabbit anti-phospho-histone3 (PH3; 1:200 Molecular Probes)-positive hemocytes were scored in randomly selected 1000 µm^2^ areas of imaged lobes.

Samples were mounted in VectaShield (Vector Laboratories). Lamellocytes were visualized by (a) high Phalloidin staining signal, (b) integrin-β (1:200, Developmental Studies Hybridoma Bank CF.6G11 [95]) expression, or (c) *msnf9-GFP* expression [89]. Representative results from twelve or more dissections from at least three independent experiments are presented, unless specified otherwise.

### Confocal imaging

Mounted samples were imaged with Zeiss laser scanning confocal microscopes LSM 510 or LSM710. For each experiment, images were scanned on the same microscope with the same software and scan settings. Images were gathered at 0.8 µm -1.5 µm and recorded at 8 bit. Laser amplifier gain and offset values were set with negative controls lacking either primary antibodies or wasp infection. Images were processed with Zeiss LSM image browser or Zen Lite 2012. Figures were assembled in Adobe Photoshop v12.0.4 and CC 2015.5 or Illustrator CC 2015.3.

---

**Fig S1.**
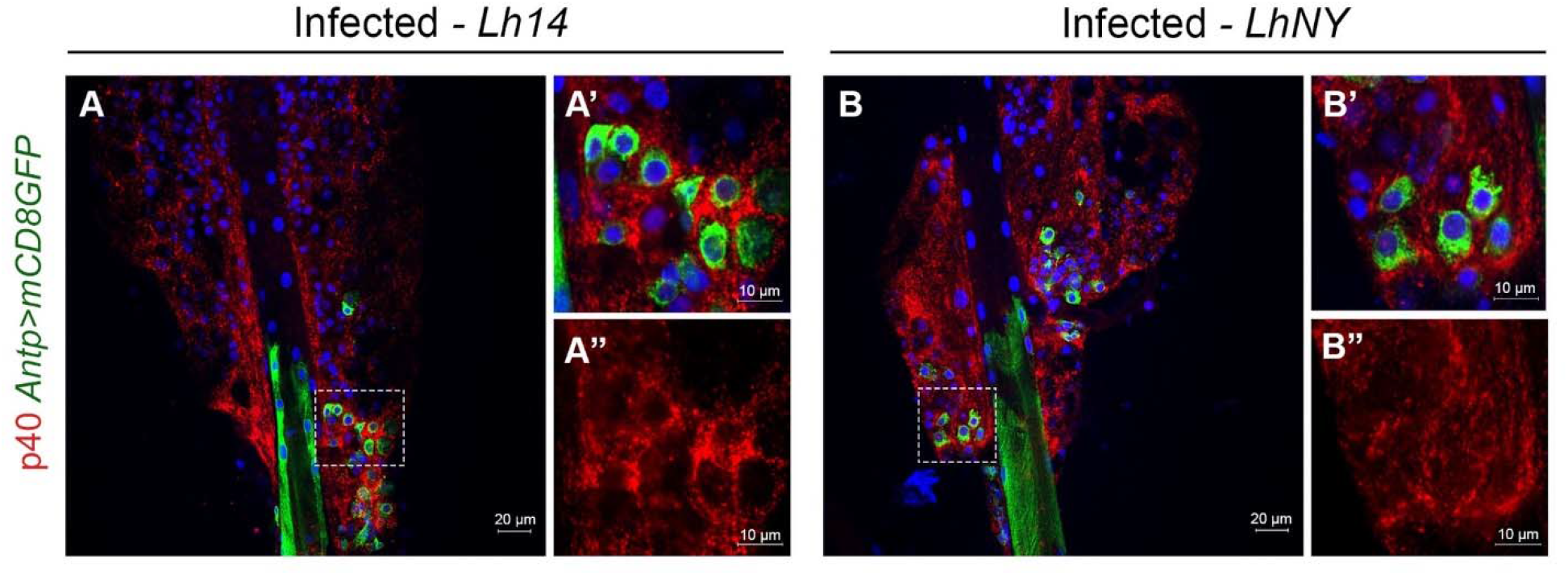
Similar effects of different *Lh* strains. (**A-B**) SSp40 staining of lymph glands from *Lh14-* **(A-A’’**) or *LhNY-*infected (**B-B”**) *Antp>mCD8GFP* hosts. Strong punctate EV signals are observed around the GFP-positive PSCs and in hemocytes. Areas in the PSC are enlarged in the insets to show details.

**Fig S2.**
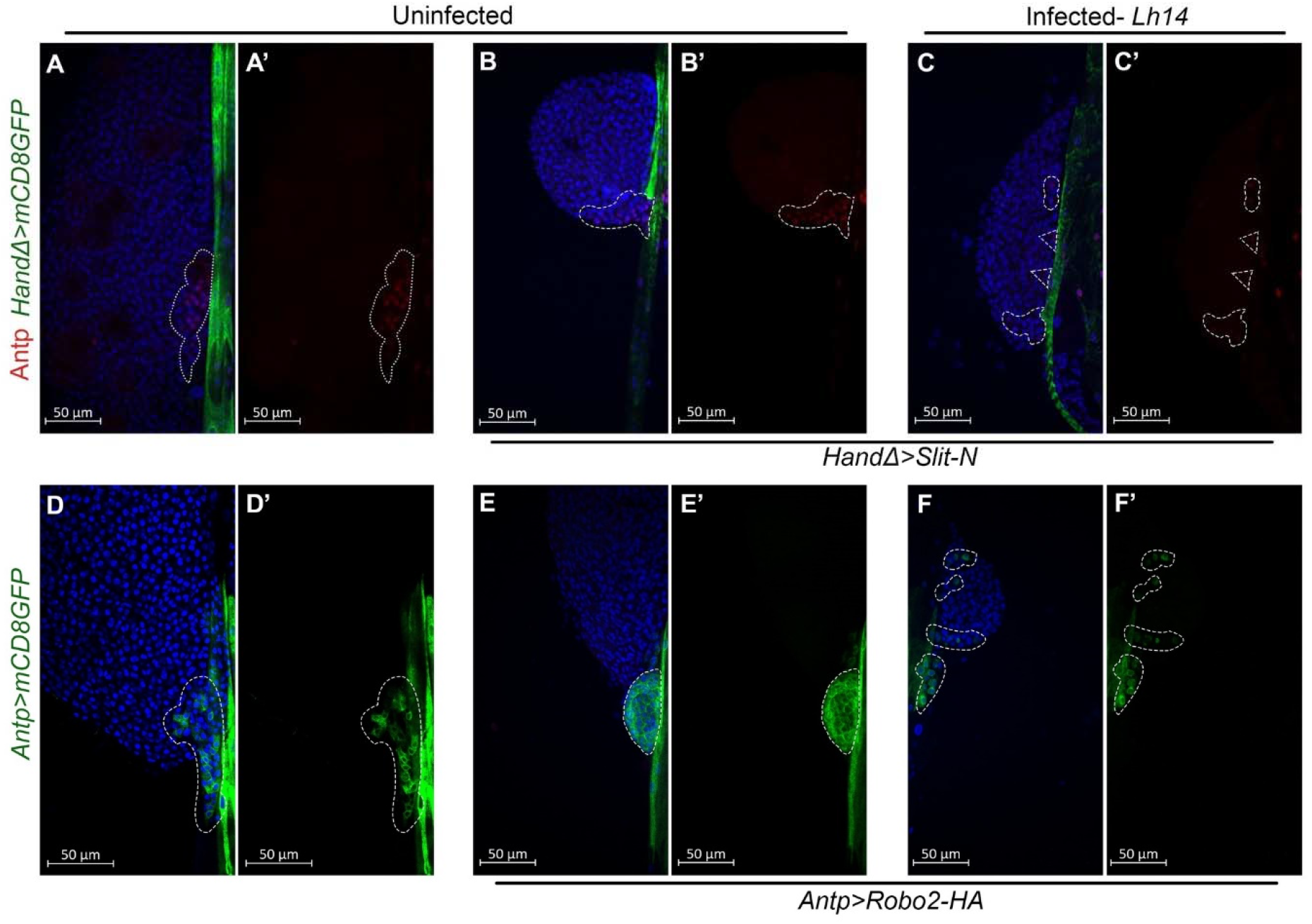
*Lh* infection overrides Slit-Robo signaling. (**A-C**) Antp staining of lymph glands from Hand*Δ>mCD8GFP* (**A-A’**) and Hand*Δ>mCD8GFP, Slit-N* (**B-C’**) hosts. The tight clustering of Antp-positive PSC in infected hosts is lost and the PSC is disassembled (**C-C’**). (**D-F**) Lymph glands from *Antp>mCD8GFP* (**D-D’**) and *Antp>mCD8GFP, Robo2-HA* hosts (**E-F’**). (**E-E’**) Robo2-HA expression tightens the GFP-positive PSC. (**F-F’**). *Lh* attack overrides this effect. *Lh* EVs are associated with these Antp> *mCD8GFP, Robo2-HA lobes* (see **Fig. S3**).

**Fig S3.**
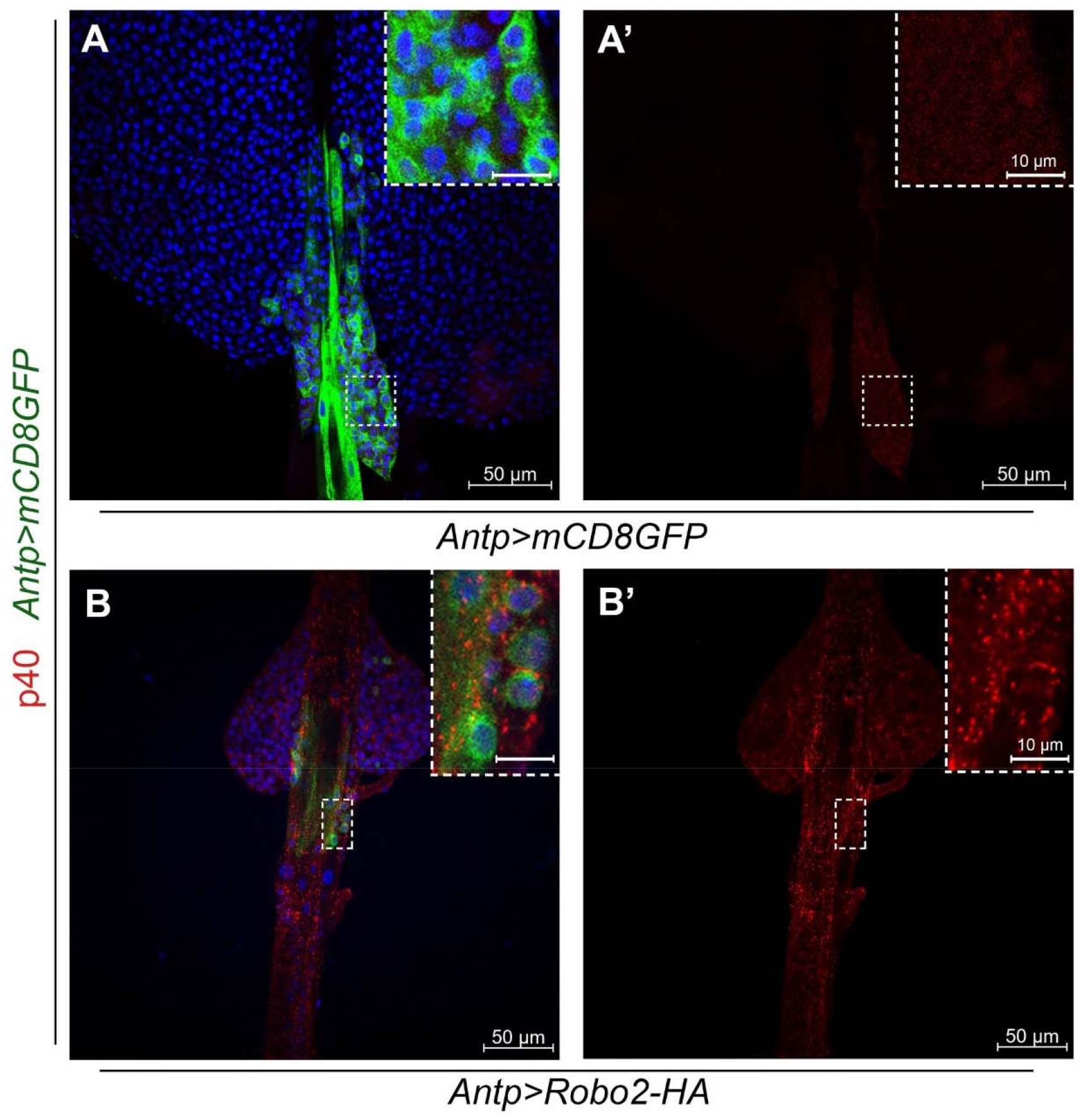
*Lh* EVs are associated with *Robo2-HA* lobes. Anterior lobes of lymph glands from uninfected *Antp>mCD8GFP* (**A-A’**) and *Antp>mCD8GFP, Robo2-HA* animals (**B-B’**). EVs are absent in glands of naïve animals (**A, A’**) but clearly observed and widely distributed in glands from infected animals. The PSC cells becoming dislodged are also evident. (The sample in panels **B, B’** is the same as shown in **Fig. S2**, panels **F, F’**.)

**Fig S4.**
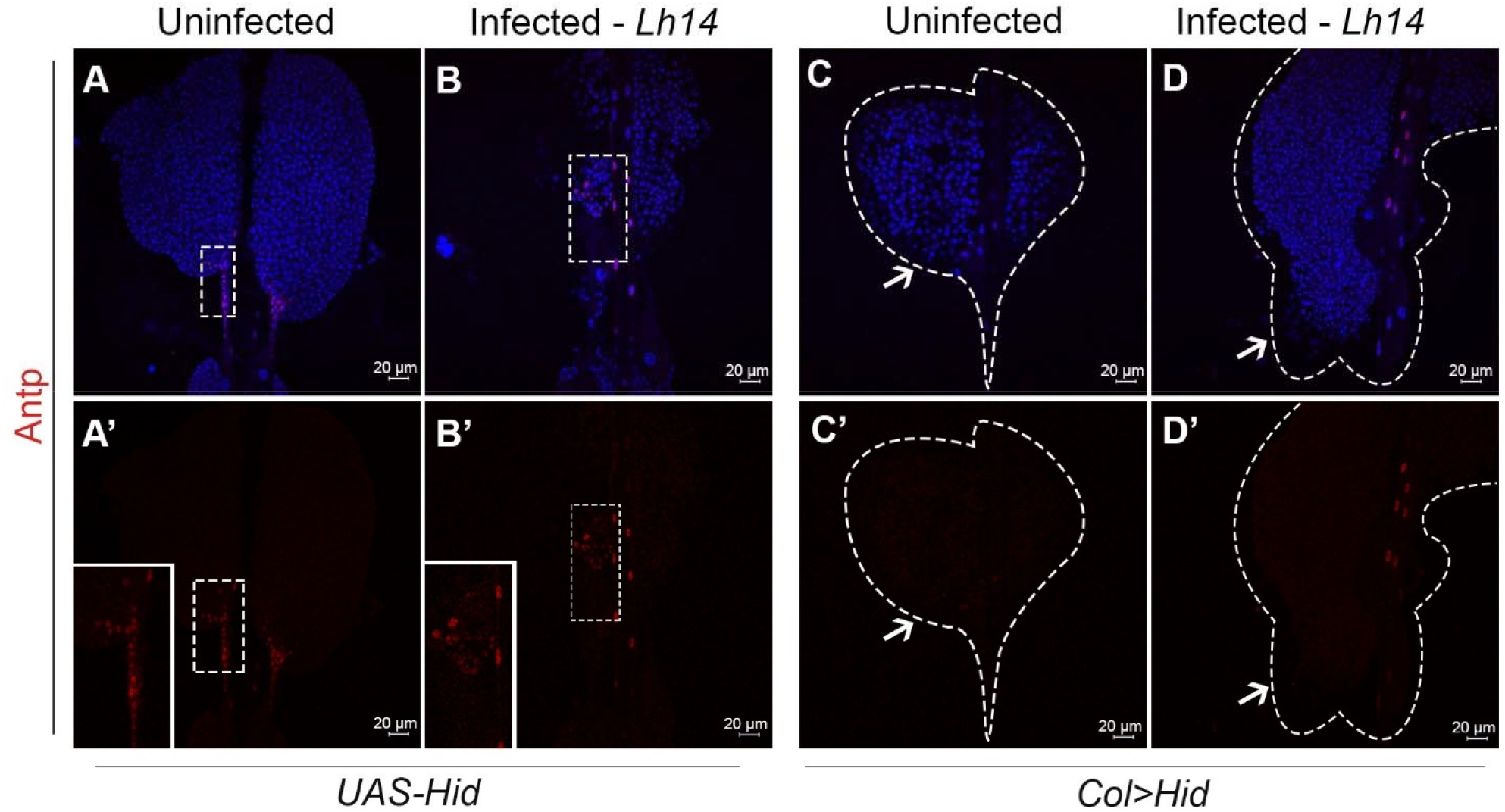
PSC-less lymph glands do not respond to *Lh* infection. (**A-A’**) A normal-sized and intact PSC, expresses Antp in *UAS-Hid* animals. Lobes from naïve animals have normal morphology. (**B-B’**) An Antp-positive PSC is disassembled in *UAS-Hid* animals after *Lh* infection. Lobes are reduced in size. Insets in panels A’ and B’ show Antp-positive PSC cells. (**C-D**) A PSC-less lymph gland from *Col>Hid* naïve and *Lh*-infected hosts. Lobes are Antp-negative. *Col>Hid* lobes remain intact after *Lh-*infection (**D-D’**). The dashed lines in panels (**C**) and (**D**) show the areas in the images where biological samples are present. Arrows point to the general locations where the PSCs should have formed.

**Fig S5.**
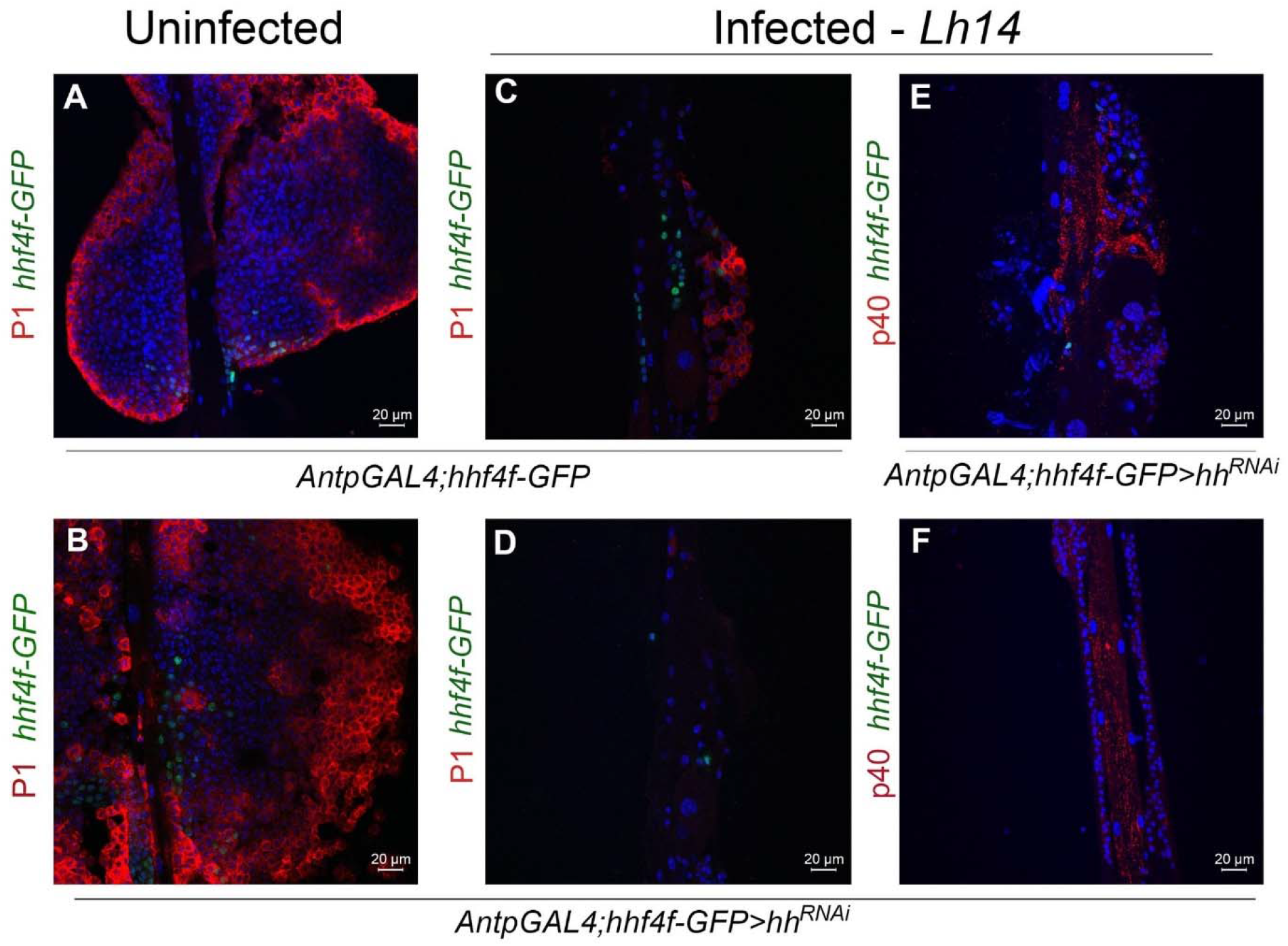
*Antp>hh*^*RNAi*^ PSCs respond to *Lh* infection. (**A, B**) Lobes from naïve *AntpGAL4; hhf4f-GFP* (**A**) and *Antp>hh*^*RNAi*^; *hhf4f-GFP* (**B**) hosts. *hh* KD increased cortical P1-positive cells. (**C, D**) Anterior lobes from *Lh-*infected hosts showing significant hemocyte loss and disassembled PSCs. In both cases, few GFP-positive PSC cells are present. P1-positive cells are also present after *Lh* infection (**C**). (**E, F**) Anterior (**E**) and posterior (**F**) lobes from *Lh-*infected *Antp>hh*^*RNAi*^; *hhf4f-GFP* hosts show many EVs in the few hemocytes remaining after infection. EVs are also evident in the dorsal vessel.

**Fig S6.**
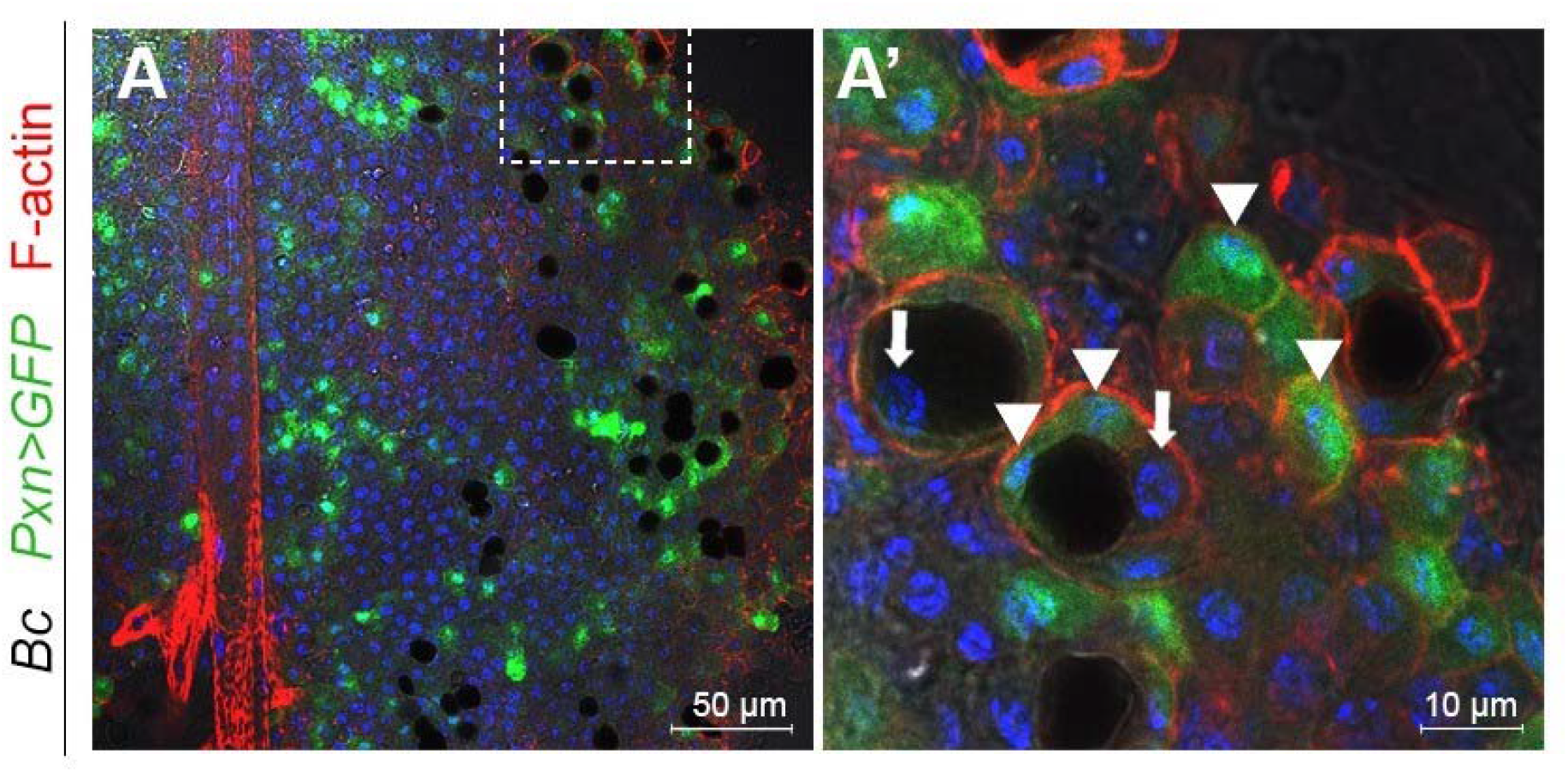
Blackened crystal cells are phagocytosed by lymph gland hemocytes. (**A A’**) A *Bc/Bc*^*+*^ *Pxn>GFP* gland showing blackened crystal cells contained within *Pxn>GFP*-expressing hemocytes. Arrows points to crystal cell nuclei; arrowheads point to *Pxn>GFP*-positive macrophages. Not all macrophages contain a crystal cell.

**Fig S7.**
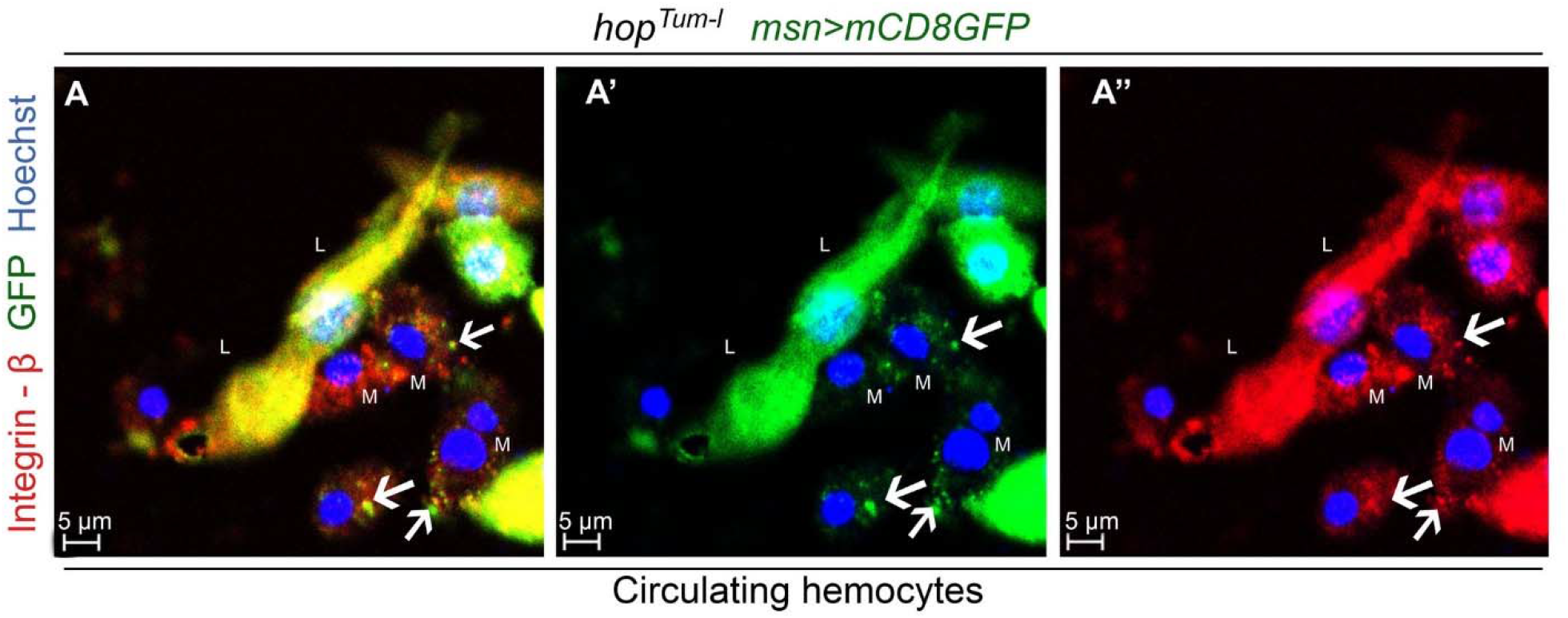
Efferocytosis of disintegrating lamellocytes. Smears from an *Lh*-infected *hop*^*Tum-l*^ host in which lamellocytes (L) express *mCD8GFP*. Lamellocytes also express high levels of integrin-beta. Double positive lamellocyte fragments in panel **A** (pseudocolors in **A’** and **A”** are merged in panel **A**) are observed in macrophages (M) indicated by arrows.

**Fig. S8.**
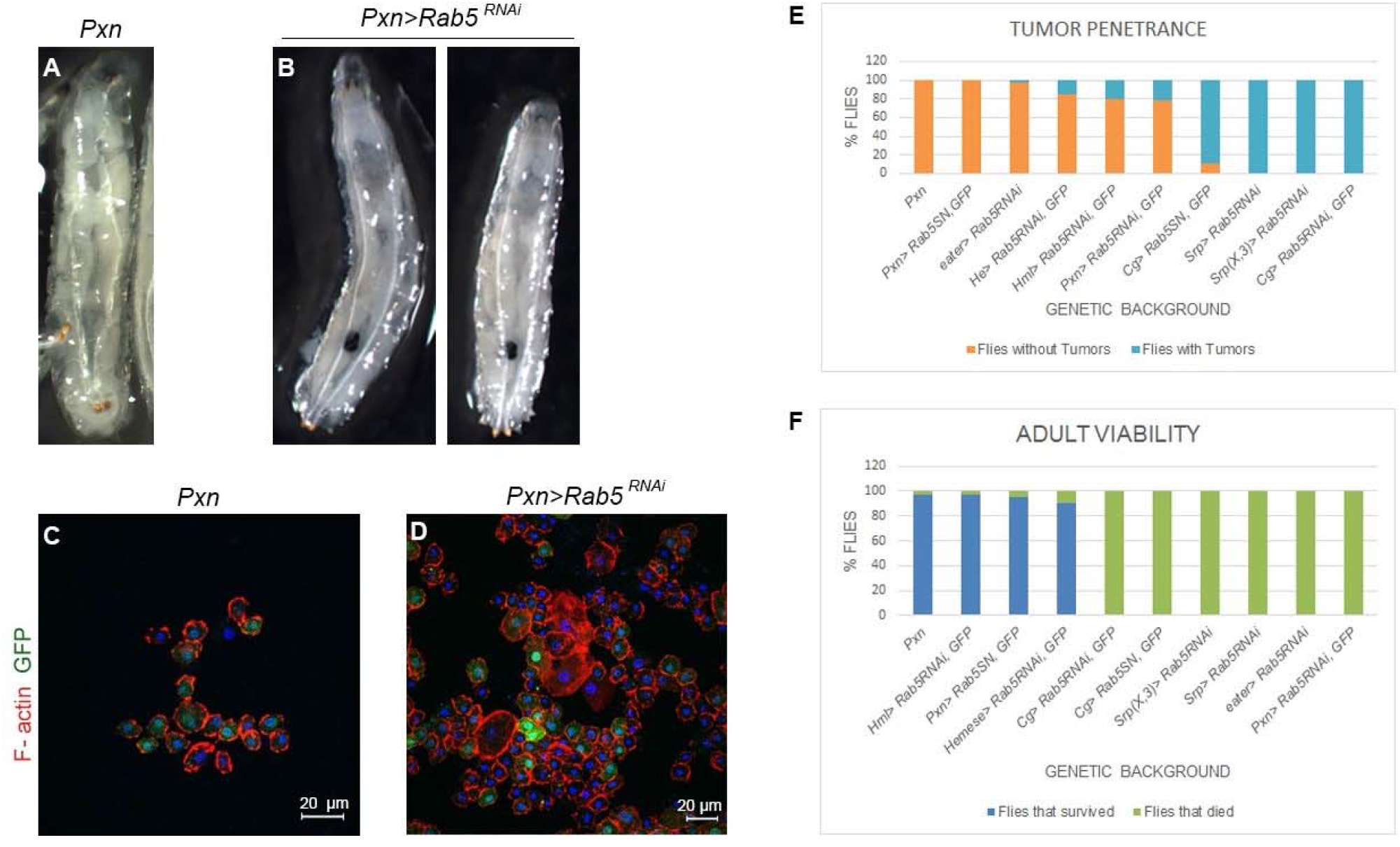
Tumorigenesis and lethality in *Rab5* knockdown animals. (**A, B**) *Pxn>GFP, UAS-Rab5*^*RNAi*^ larvae with melanized tumors clearly visible through the transparent cuticle. Tumors are absent in the control animal. (**C, D**) Smears from circulating hemocytes of these animals. An overabundance of *Pxn>GFP-*positive and GFP-negative (lamellocytes) cells is observed. (**E**) Tumor penetrance (animals with tumors/animals scored) in *Rab5* KD animals varied with different GAL4 drivers. (**F**) Viability to adulthood was differentially compromised. More than 100 animals were scored for each cross in panels (**E**) and (**F**).

## Notes

### Competing Interest Statement

The authors have declared no competing interest.

## Literature cited

1. Libersat F, Delago A, Gal R. Manipulation of host behavior by parasitic insects and insect parasites. Annu Rev Entomol. 2009;54:189-207. Epub 2008/12/11. doi: 10.1146/annurev.ento.54.110807.090556. PubMed PMID: 19067631.

2. Pennacchio F, Strand, M.R. Evolution of developmental strategies in parasitic Hymenoptera. Annu Rev Entomol. 2006;51:233-58.

3. Poirie M, Carton Y, Dubuffet, A. Virulence strategies in parasitoid Hymenoptera as an example of adaptive diversity. C R Biol. 2009;332(2-3):311-20. Epub 2009/03/14. doi: 10.1016/j.crvi.2008.09.004. PubMed PMID: 19281961.

4. Keebaugh ES, Schlenke, TA. Insights from natural host-parasite interactions: the Drosophila model. Dev Comp Immunol. 2014;42(1):111-23. Epub 2013/06/15. doi: 10.1016/j.dci.2013.06.001. PubMed PMID: 23764256; PubMed Central PMCID: PMCPMC3808516.

5. Heavner ME, Hudgins AD, Rajwani R, Morales J, Govind, S. Harnessing the natural -parasitoid model for integrating insect immunity with functional venomics. Curr Opin Insect Sci. 2014;6:61-7. Epub 2015/02/03. doi: 10.1016/j.cois.2014.09.016. PubMed PMID: 25642411; PubMed Central PMCID: PMC4309977.

6. Small C, Paddibhatla I, Rajwani R, Govind, S. An introduction to parasitic wasps of Drosophila and the antiparasite immune response. J Vis Exp. 2012;(63):e3347. Epub 2012/05/17. doi: 3347 [pii] 10.3791/3347. PubMed PMID: 22588641.

7. Schlenke TA, Morales J, Govind S, Clark, AG. Contrasting infection strategies in generalist and specialist wasp parasitoids of Drosophila melanogaster. PLoS Pathog. 2007;3(10):1486-501. Epub 2007/10/31. doi: 06-PLPA-RA-0488 [pii] 10.1371/journal.ppat.0030158. PubMed PMID: 17967061; PubMed Central PMCID: PMC2042021.

8. Sorrentino RP, Carton Y, Govind, S. Cellular immune response to parasite infection in the Drosophila lymph gland is developmentally regulated. Dev Biol. 2002;243(1):65-80. Epub 2002/02/16. doi: 10.1006/dbio.2001.0542 S0012160601905421 [pii]. PubMed PMID: 11846478.

9. Lanot R, Zachary D, Holder F, Meister, M. Postembryonic hematopoiesis in Drosophila. Dev Biol. 2001;230(2):243-57. Epub 2001/02/13. doi: 10.1006/dbio.2000.0123 S0012-1606(00)90123-4 [pii]. PubMed PMID: 11161576.

10. Anderl I, Vesala L, Ihalainen TO, Vanha-aho L-M, Andó I, Rämet M, et, al. Transdifferentiation and proliferation in two distinct hemocyte lineages in drosophila melanogaster larvae after wasp infection. PLoS Pathog. 2016;12(7):e1005746. doi: 10.1371/journal.ppat.1005746. PubMed PMID: PMC4945071.

11. Huang J, Chen J, Fang G, Pang L, Zhou S, Zhou Y, et, al. Two novel venom proteins underlie divergent parasitic strategies between a generalist and a specialist parasite. Nat Commun. 2021;12(1):234. Epub 2021/01/13. doi: 10.1038/s41467-020-20332-8. PubMed PMID: 33431897; PubMed Central PMCID: PMCPMC7801585.

12. Rizki TM, Rizki, R.M., Carton, Y. Leptopilina heterotoma and L. boulardi: strategies to avoid cellular defense responses of Drosophila melanogaster. Exp Parasitol. 1990;70:466-75.

13. Chiu H, Govind, S. Natural infection of, D. melanogaster by virulent parasitic wasps induces apoptotic depletion of hematopoietic precursors. Cell Death Differ. 2002;9(12):1379-81. Epub 2002/12/13. doi: 10.1038/sj.cdd.4401134. PubMed PMID: 12478476.

14. Wan B, Goguet E, Ravallec M, Pierre O, Lemauf S, Volkoff AN, et, al. Venom Atypical Extracellular Vesicles as Interspecies Vehicles of Virulence Factors Involved in Host Specificity: The Case of a Drosophila Parasitoid Wasp. Front Immunol. 2019;10:1688. Epub 2019/08/06. doi: 10.3389/fimmu.2019.01688. PubMed PMID: 31379874; PubMed Central PMCID: PMCPMC6653201.

15. Heavner ME, Ramroop J, Gueguen G, Ramrattan G, Dolios G, Scarpati M, et, al. Novel Organelles with Elements of Bacterial and Eukaryotic Secretion Systems Weaponize Parasites of Drosophila. Curr Biol. 2017;27(18):2869-77 e6. Epub 2017/09/12. doi: 10.1016/j.cub.2017.08.019. PubMed PMID: 28889977; PubMed Central PMCID: PMCPMC5659752.

16. Rizki RM, Rizki, TM. Parasitoid virus-like particles destroy Drosophila cellular immunity. Proc Natl Acad Sci U S A. 1990;87(21):8388-92. Epub 1990/11/01. PubMed PMID: 2122461; PubMed Central PMCID: PMC54961.

17. Dupas S, Brehelin M, Frey F, Carton, Y. Immune suppressive virus-like particles in a Drosophila parasitoid: significance of their intraspecific morphological variations. Parasitology. 1996;113 (Pt 3):207- 12. Epub 1996/09/01. PubMed PMID: 8811846.

18. Morales J, Chiu H, Oo T, Plaza R, Hoskins S, Govind, S. Biogenesis, structure, and immune- suppressive effects of virus-like particles of a Drosophila parasitoid, Leptopilina victoriae. J Insect Physiol. 2005;51(2):181-95. Epub 2005/03/08. doi: S0022-1910(04)00189-1 [pii] 10.1016/j.jinsphys.2004.11.002. PubMed PMID: 15749103.

19. Di Giovanni D, Lepetit D, Guinet B, Bennetot B, Boulesteix M, Coute Y, et, al. A Behavior- Manipulating Virus Relative as a Source of Adaptive Genes for Drosophila Parasitoids. Mol Biol Evol. 2020;37(10):2791-807. Epub 2020/02/23. doi: 10.1093/molbev/msaa030. PubMed PMID: 32080746.

20. Chiu H, Morales J, Govind, S. Identification and immuno-electron microscopy localization of p40, a protein component of immunosuppressive virus-like particles from Leptopilina heterotoma, a virulent parasitoid wasp of Drosophila. J Gen Virol. 2006;87(Pt 2):461-70. Epub 2006/01/25. doi: 87/2/461 [pii] 10.1099/vir.0.81474-0. PubMed PMID: 16432035; PubMed Central PMCID: PMC2705942.

21. Gueguen G, Rajwani R, Paddibhatla I, Morales J, Govind, S. VLPs of Leptopilina boulardi share biogenesis and overall stellate morphology with VLPs of the heterotoma clade. Virus Res. 2011;160(1- 2):159-65. Epub 2011/06/28. doi: S0168-1702(11)00228-0 [pii] 10.1016/j.virusres.2011.06.005. PubMed PMID: 21704090.

22. Ferrarese R, Morales J, Fimiarz D, Webb BA, Govind, S. A supracellular system of actin-lined canals controls biogenesis and release of virulence factors in parasitoid venom glands. J Exp Biol. 2009;212(Pt 14):2261-8. Epub 2009/06/30. doi: 212/14/2261 [pii] 10.1242/jeb.025718. PubMed PMID: 19561216; PubMed Central PMCID: PMC2702457.

23. Wey B, Heavner ME, Wittmeyer KT, Briese T, Hopper KR, Govind, S. Immune Suppressive Extracellular Vesicle Proteins of Leptopilina heterotoma Are Encoded in the Wasp Genome. G3 (Bethesda). 2020;10(1):1-12. Epub 2019/11/05. doi: 10.1534/g3.119.400349. PubMed PMID: 31676506; PubMed Central PMCID: PMCPMC6945029.

24. Espina M, Olive AJ, Kenjale R, Moore DS, Ausar SF, Kaminski RW, et, al. IpaD localizes to the tip of the type III secretion system needle of Shigella flexneri. Infect Immun. 2006;74(8):4391-400. Epub 2006/07/25. doi: 74/8/4391 [pii] 10.1128/IAI.00440-06. PubMed PMID: 16861624; PubMed Central PMCID: PMC1539624.

25. Arizmendi O, Picking, W. D., and Picking, W. L. Macrophage Apoptosis Triggered by IpaD from Shigella flexneri. Infection and Immunity. 2016;84(6):1857-65. doi: 10.1128/IAI.01483-15.

26. Colinet D, Schmitz, A., Cazes, D., Gatti, J.-L., Poirie, M. The origin of intraspecific variation of virulence in an eukaryotic immune suppressive parasite. PLoS Pathog. 2010;6:e1001206.

27. Stofanko M, Kwon, S.Y., Badenhorst, P. Lineage tracing of lamellocytes demonstrates Drosophila macrophage plasticity. PLoS ONE. 2010;5:e14051. doi: doi: 10.1371/journal.pone.0014051.

28. Rotstein B, Paululat, A. On the Morphology of the Drosophila Heart. J Cardiovasc Dev Dis. 2016;3(2). Epub 2016/04/12. doi: 10.3390/jcdd3020015. PubMed PMID: 29367564; PubMed Central PMCID: PMCPMC5715677.

29. Rizki, T. The circulatory system and associated cells and tissues. In: Ashburner MaWT, editor. The Genetics and Biology of Drosophila. 2b. London: Academic Press; 1978. p. 397-452.

30. Banerjee U, Girard JR, Goins LM, Spratford, CM. Drosophila as a Genetic Model for Hematopoiesis. Genetics. 2019;211(2):367-417. Epub 2019/02/09. doi: 10.1534/genetics.118.300223. PubMed PMID: 30733377; PubMed Central PMCID: PMCPMC6366919.

31. Mandal L, Martinez-Agosto JA, Evans CJ, Hartenstein V, Banerjee, U. A Hedgehog- and Antennapedia-dependent niche maintains Drosophila haematopoietic precursors. Nature. 2007;446(7133):320-4. Epub 2007/03/16. doi: nature05585 [pii] 10.1038/nature05585. PubMed PMID: 17361183; PubMed Central PMCID: PMC2807630.

32. Krzemien J, Dubois L, Makki R, Meister M, Vincent A, Crozatier, M. Control of blood cell homeostasis in Drosophila larvae by the posterior signalling centre. Nature. 2007;446(7133):325-8. Epub 2007/03/16. doi: nature05650 [pii] 10.1038/nature05650. PubMed PMID: 17361184.

33. Crozatier M, Glise B, Vincent, A. Patterns in evolution: veins of the Drosophila wing. Trends Genet. 2004;20(10):498-505. Epub 2004/09/15. doi: 10.1016/j.tig.2004.07.013 S0168-9525(04)00221-5 [pii]. PubMed PMID: 15363904.

34. Krzemien J, Crozatier M, Vincent, A. Ontogeny of the Drosophila larval hematopoietic organ, hemocyte homeostasis and the dedicated cellular immune response to parasitism. Int J Dev Biol. 2010;54(6-7):1117-25. Epub 2010/08/17. doi: 093053jk [pii] 10.1387/ijdb.093053jk. PubMed PMID: 20711989.

35. Gueguen G, Kalamarz, M.E., Ramroop, J., Uribe, J., Govind, S. Polydnaviral ankyrin proteins aid parasitic wasp survival by coordinate and selective inhibition of hematopoietic and immune NF-kappa B signaling in insect hosts. PLoS Pathog. 2013;9 (8):e1003580.

36. Benmimoun B, Polesello C, Haenlin M, Waltzer, L. The EBF transcription factor Collier directly promotes Drosophila blood cell progenitor maintenance independently of the niche. Proc Natl Acad Sci U S A. 2015;112(29):9052-7. Epub 2015/07/08. doi: 1423967112 [pii] 10.1073/pnas.1423967112. PubMed PMID: 26150488; PubMed Central PMCID: PMC4517242.

37. Sinenko SA, Shim J, Banerjee, U. Oxidative stress in the haematopoietic niche regulates the cellular immune response in Drosophila. EMBO Rep. 2012;13(1):83-9. Epub 2011/12/03. doi: embor2011223 [pii] 10.1038/embor.2011.223. PubMed PMID: 22134547; PubMed Central PMCID: PMC3246251.

38. Small C, Ramroop J, Otazo M, Huang LH, Saleque S, Govind, S. An unexpected link between notch signaling and ROS in restricting the differentiation of hematopoietic progenitors in Drosophila. Genetics. 2014;197(2):471-83. Epub 2013/12/10. doi: genetics.113.159210 [pii] 10.1534/genetics.113.159210. PubMed PMID: 24318532; PubMed Central PMCID: PMC4063908.

39. Khadilkar RJ, Vogl W, Goodwin K, Tanentzapf, G. Modulation of occluding junctions alters the hematopoietic niche to trigger immune activation. Elife. 2017;6. Epub 2017/08/26. doi: 10.7554/eLife.28081. PubMed PMID: 28841136; PubMed Central PMCID: PMCPMC5597334.

40. Grigorian M, Mandal L, Hartenstein, V. Hematopoiesis at the onset of metamorphosis: terminal differentiation and dissociation of the Drosophila lymph gland. Dev Genes Evol. 2011;221(3):121-31. Epub 2011/04/22. doi: 10.1007/s00427-011-0364-6. PubMed PMID: 21509534.

41. Morin X, Daneman R, Zavortink M, Chia, W. A protein trap strategy to detect GFP-tagged proteins expressed from their endogenous loci in Drosophila. Proc Natl Acad Sci U S A. 2001;98(26):15050-5. Epub 2001/12/14. doi: 10.1073/pnas.261408198 261408198 [pii]. PubMed PMID: 11742088; PubMed Central PMCID: PMC64981.

42. Kelso RJ, Buszczak M, Quinones AT, Castiblanco C, Mazzalupo S, Cooley, L. Flytrap, a database documenting a GFP protein-trap insertion screen in Drosophila melanogaster. Nucleic Acids Res. 2004;32(Database issue):D418-20. Epub 2003/12/19. doi: 10.1093/nar/gkh014. PubMed PMID: 14681446; PubMed Central PMCID: PMCPMC308749.

43. Crozatier M, Ubeda JM, Vincent A, Meister, M. Cellular immune response to parasitization in Drosophila requires the EBF orthologue collier. PLoS Biol. 2004;2(8):E196. Epub 2004/08/18. doi: 10.1371/journal.pbio.0020196. PubMed PMID: 15314643; PubMed Central PMCID: PMC509289.

44. Louradour I, Sharma A, Morin-Poulard I, Letourneau M, Vincent A, Crozatier M, et, al. Reactive oxygen species-dependent Toll/NF-kappaB activation in the Drosophila hematopoietic niche confers resistance to wasp parasitism. Elife. 2017;6. Epub 2017/11/02. doi: 10.7554/eLife.25496. PubMed PMID: 29091025; PubMed Central PMCID: PMCPMC5681226.

45. Morin-Poulard I, Sharma A, Louradour I, Vanzo N, Vincent A, Crozatier, M. Vascular control of the Drosophila haematopoietic microenvironment by Slit/Robo signalling. Nat Commun. 2016;7:11634. Epub 2016/05/20. doi: 10.1038/ncomms11634. PubMed PMID: 27193394; PubMed Central PMCID: PMCPMC4874035.

46. Ordan E, Brankatschk M, Dickson B, Schnorrer F, Volk, T. Slit cleavage is essential for producing an active, stable, non-diffusible short-range signal that guides muscle migration. Development. 2015;142(8):1431-6. Epub 2015/03/31. doi: 10.1242/dev.119131. PubMed PMID: 25813540.

47. Bhuin T, Roy, JK. Rab proteins: the key regulators of intracellular vesicle transport. Exp Cell Res. 2014;328(1):1-19. Epub 2014/08/05. doi: S0014-4827(14)00318-8 [pii] 10.1016/j.yexcr.2014.07.027. PubMed PMID: 25088255.

48. Zhang J, Schulze KL, Hiesinger PR, Suyama K, Wang S, Fish M, et, al. Thirty-one flavors of Drosophila rab proteins. Genetics. 2007;176(2):1307-22. Epub 2007/04/06. doi: 10.1534/genetics.106.066761. PubMed PMID: 17409086; PubMed Central PMCID: PMCPMC1894592.

49. Rizki TM, Rizki, RM. Parasitoid-induced cellular immune deficiency in Drosophila. Ann N Y Acad Sci. 1994;712:178-94. Epub 1994/04/15. PubMed PMID: 7910721.

50. Melk JP, Govind, S. Developmental analysis of Ganaspis xanthopoda, a larval parasitoid of Drosophila melanogaster. J Exp Biol. 1999;202(Pt 14):1885-96. Epub 1999/06/23. PubMed PMID: 10377270.

51. Chiu H, Sorrentino RP, Govind, S. Suppression of the Drosophila cellular immune response by Ganaspis xanthopoda. Adv Exp Med Biol. 2001;484:161-7. Epub 2001/06/23. doi: 10.1007/978-1-4615-1291-2_14. PubMed PMID: 11418981.

52. Suzuki M, Tanaka, T. Virus-like particles in venom of Meteorus pulchricornis induce host hemocyte apoptosis. J Insect Physiol. 2006;52(6):602-13. Epub 2006/05/23. doi: 10.1016/j.jinsphys.2006.02.009. PubMed PMID: 16712867.

53. Teramoto T, Tanaka, T. Mechanism of reduction in the number of the circulating hemocytes in the Pseudaletia separata host parasitized by Cotesia kariyai. J Insect Physiol. 2004;50(12):1103-11. Epub 2005/01/27. doi: 10.1016/j.jinsphys.2004.08.005. PubMed PMID: 15670857.

54. Wan NF, Ji XY, Zhang H, Yang JH, Jiang, JX. Nucleopolyhedrovirus infection and/or parasitism by Microplitis pallidipes Szepligeti affect hemocyte apoptosis of Spodoptera exigua (Hubner) larvae. J Invertebr Pathol. 2015;132:165-70. Epub 2015/10/17. doi: 10.1016/j.jip.2015.10.004. PubMed PMID: 26470677.

55. Jung SH, Evans CJ, Uemura C, Banerjee, U. The Drosophila lymph gland as a developmental model of hematopoiesis. Development. 2005;132(11):2521-33. Epub 2005/04/29. doi: dev.01837 [pii] 10.1242/dev.01837. PubMed PMID: 15857916.

56. Kim-Jo C, Gatti JL, Poirie, M. Drosophila Cellular Immunity Against Parasitoid Wasps: A Complex and Time-Dependent Process. Front Physiol. 2019;10:603. Epub 2019/06/04. doi: 10.3389/fphys.2019.00603. PubMed PMID: 31156469; PubMed Central PMCID: PMCPMC6529592.

57. Baldeosingh R, Gao H, Wu X, Fossett, N. Hedgehog signaling from the Posterior Signaling Center maintains U-shaped expression and a prohemocyte population in Drosophila. Dev Biol. 2018;441(1):132- 45. Epub 2018/07/04. doi: 10.1016/j.ydbio.2018.06.020. PubMed PMID: 29966604; PubMed Central PMCID: PMCPMC6064674.

58. Tokusumi Y, Tokusumi T, Shoue DA, Schulz, RA. Gene regulatory networks controlling hematopoietic progenitor niche cell production and differentiation in the Drosophila lymph gland. PLoS One. 2012;7(7):e41604. Epub 2012/08/23. doi: 10.1371/journal.pone.0041604 PONE-D-12-16104 [pii]. PubMed PMID: 22911822; PubMed Central PMCID: PMC3404040.

59. Sinenko SA, Mandal L, Martinez-Agosto JA, Banerjee, U. Dual role of wingless signaling in stem-like hematopoietic precursor maintenance in Drosophila. Dev Cell. 2009;16(5):756-63. Epub 2009/05/23. doi: S1534-5807(09)00093-8 [pii] 10.1016/j.devcel.2009.03.003. PubMed PMID: 19460351; PubMed Central PMCID: PMC2718753.

60. Krzemien J, Oyallon J, Crozatier M, Vincent, A. Hematopoietic progenitors and hemocyte lineages in the Drosophila lymph gland. Dev Biol. 2010;346(2):310-9. Epub 2010/08/17. doi: S0012-1606(10)00987-5 [pii] 10.1016/j.ydbio.2010.08.003. PubMed PMID: 20707995.

61. Benmimoun B, Haenlin M, Waltzer, L. Haematopoietic progenitor maintenance by EBF/Collier: beyond the Niche. Cell Cycle. 2015;14(22):3517-8. Epub 2015/12/15. doi: 10.1080/15384101.2015.1093449. PubMed PMID: 26654595.

62. Sung BH, Weaver, AM. Exosome secretion promotes chemotaxis of cancer cells. Cell Adh Migr. 2017;11(2):187-95. Epub 2017/01/28. doi: 10.1080/19336918.2016.1273307. PubMed PMID: 28129015; PubMed Central PMCID: PMCPMC5351719.

63. Majumdar R, Tavakoli Tameh A, Parent, CA. Exosomes Mediate LTB4 Release during Neutrophil Chemotaxis. PLoS Biol. 2016;14(1):e1002336. Epub 2016/01/08. doi: 10.1371/journal.pbio.1002336. PubMed PMID: 26741884; PubMed Central PMCID: PMCPMC4704783.

64. Case EDR, Samuel, JE. Contrasting Lifestyles Within the Host Cell. Microbiol Spectr. 2016;4(1). Epub 2016/03/22. doi: 10.1128/microbiolspec.VMBF-0014-2015. PubMed PMID: 26999394; PubMed Central PMCID: PMCPMC4804636.

65. Heavner, M. Evidence for organelle-like extracellular vesicles from a parasite of Drosophila and their fundion in suppressing host immunity. https://academicworkscunyedu/gc_etds/2585. 2018.

66. Zhang Y, Bliska, JB. Role of macrophage apoptosis in the pathogenesis of Yersinia. Curr Top Microbiol Immunol. 2005;289:151-73. Epub 2005/03/29. PubMed PMID: 15791955.

67. Hueffer K, Galan, JE. Salmonella-induced macrophage death: multiple mechanisms, different outcomes. Cell Microbiol. 2004;6(11):1019-25. Epub 2004/10/08. doi: 10.1111/j.1462-5822.2004.00451.x. PubMed PMID: 15469431.

68. Jarvelainen HA, Galmiche A, Zychlinsky, A. Caspase-1 activation by Salmonella. Trends Cell Biol. 2003;13(4):204-9. Epub 2003/04/02. PubMed PMID: 12667758.

69. Wan B, Poirie M, Gatti, JL. Parasitoid wasp venom vesicles (venosomes) enter Drosophila melanogaster lamellocytes through a flotillin/lipid raft-dependent endocytic pathway. Virulence. 2020;11(1):1512-21. Epub 2020/11/03. doi: 10.1080/21505594.2020.1838116. PubMed PMID: 33135553; PubMed Central PMCID: PMCPMC7605353.

70. Del Signore SJ, Biber SA, Lehmann KS, Heimler SR, Rosenfeld BH, Eskin TL, et, al. dOCRL maintains immune cell quiescence by regulating endosomal traffic. PLoS Genet. 2017;13(10):e1007052. Epub 2017/10/14. doi: 10.1371/journal.pgen.1007052. PubMed PMID: 29028801; PubMed Central PMCID: PMCPMC5656325.

71. Shravage BV, Hill JH, Powers CM, Wu L, Baehrecke, EH. Atg6 is required for multiple vesicle trafficking pathways and hematopoiesis in Drosophila. Development. 2013;140(6):1321-9. Epub 2013/02/15. doi: 10.1242/dev.089490. PubMed PMID: 23406899; PubMed Central PMCID: PMCPMC3585664.

72. Zhou B, Yun, E.Y., Ray, L., You, J., Ip, Y.T., Lin, X. Retromer promotes immune quiescence by suppressing Spätzle-Toll pathway in Drosophila. J Cell Physiology. 2014;229:512-20.

73. Emerald BS, Cohen, SM. Spatial and temporal regulation of the homeotic selector gene Antennapedia is required for the establishment of leg identity in Drosophila. Dev Biol. 2004;267(2):462- 72. Epub 2004/03/12. doi: 10.1016/j.ydbio.2003.12.006 S0012160603007863 [pii]. PubMed PMID: 15013806.

74. Popichenko D, Sellin J, Bartkuhn M, Paululat, A. Hand is a direct target of the forkhead transcription factor Biniou during Drosophila visceral mesoderm differentiation. BMC Dev Biol. 2007;7:49. Epub 2007/05/22. doi: 10.1186/1471-213X-7-49. PubMed PMID: 17511863; PubMed Central PMCID: PMCPMC1891290.

75. Stramer B, Wood W, Galko MJ, Redd MJ, Jacinto A, Parkhurst SM, et, al. Live imaging of wound inflammation in Drosophila embryos reveals key roles for small GTPases during in vivo cell migration. J Cell Biol. 2005;168(4):567-73. Epub 2005/02/09. doi: 10.1083/jcb.200405120. PubMed PMID: 15699212; PubMed Central PMCID: PMCPMC2171743.

76. Kurucz E, Zettervall CJ, Sinka R, Vilmos P, Pivarcsi A, Ekengren S, et, al. Hemese, a hemocyte- specific transmembrane protein, affects the cellular immune response in Drosophila. Proc Natl Acad Sci U S A. 2003;100(5):2622-7. Epub 2003/02/25. doi: 10.1073/pnas.0436940100 0436940100 [pii]. PubMed PMID: 12598653; PubMed Central PMCID: PMC151390.

77. Tokusumi T, Shoue DA, Tokusumi Y, Stoller JR, Schulz, RA. New hemocyte-specific enhancer- reporter transgenes for the analysis of hematopoiesis in Drosophila. Genesis. 2009;47(11):771-4. Epub 2009/10/16. doi: 10.1002/dvg.20561. PubMed PMID: 19830816.

78. Asha H, Nagy I, Kovacs G, Stetson D, Ando I, Dearolf, CR. Analysis of Ras-induced overproliferation in Drosophila hemocytes. Genetics. 2003;163(1):203-15. Epub 2003/02/15. PubMed PMID: 12586708; PubMed Central PMCID: PMC1462399.

79. Waltzer L, Ferjoux G, Bataille L, Haenlin, M. Cooperation between the GATA and RUNX factors Serpent and Lozenge during Drosophila hematopoiesis. EMBO J. 2003;22(24):6516-25. Epub 2003/12/06. doi: 10.1093/emboj/cdg622. PubMed PMID: 14657024; PubMed Central PMCID: PMCPMC291817.

80. Kroeger PT, Jr., Tokusumi T, Schulz, RA. Transcriptional regulation of eater gene expression in Drosophila blood cells. Genesis. 2012;50(1):41-9. Epub 2011/08/03. doi: 10.1002/dvg.20787. PubMed PMID: 21809435.

81. Goto A, Kadowaki T, Kitagawa, Y. Drosophila hemolectin gene is expressed in embryonic and larval hemocytes and its knock down causes bleeding defects. Dev Biol. 2003;264(2):582-91. Epub 2003/12/04. doi: S0012160603004639 [pii]. PubMed PMID: 14651939.

82. Spitzweck B, Brankatschk M, Dickson, BJ. Distinct protein domains and expression patterns confer divergent axon guidance functions for Drosophila Robo receptors. Cell. 2010;140(3):409-20. Epub 2010/02/11. doi: 10.1016/j.cell.2010.01.002. PubMed PMID: 20144763.

83. Wucherpfennig T, Wilsch-Brauninger M, Gonzalez-Gaitan, M. Role of Drosophila Rab5 during endosomal trafficking at the synapse and evoked neurotransmitter release. J Cell Biol. 2003;161(3):609- 24. doi: 10.1083/jcb.200211087. PubMed PMID: 12743108; PubMed Central PMCID: PMCPMC2172938.

84. Pulipparacharuvil S, Akbar MA, Ray S, Sevrioukov EA, Haberman AS, Rohrer J, et, al. Drosophila Vps16A is required for trafficking to lysosomes and biogenesis of pigment granules. J Cell Sci. 2005;118(Pt 16):3663-73. doi: 10.1242/jcs.02502. PubMed PMID: 16046475.

85. Perkins LA, Holderbaum L, Tao R, Hu Y, Sopko R, McCall K, et, al. The Transgenic RNAi Project at Harvard Medical School: Resources and Validation. Genetics. 2015;201(3):843-52. Epub 2015/09/01. doi: 10.1534/genetics.115.180208. PubMed PMID: 26320097; PubMed Central PMCID: PMCPMC4649654.

86. Lemaitre B, Kromer-Metzger E, Michaut L, Nicolas E, Meister M, Georgel P, et, al. A recessive mutation, immune deficiency (imd), defines two distinct control pathways in the Drosophila host defense. Proc Natl Acad Sci U S A. 1995;92(21):9465-9. Epub 1995/10/10. PubMed PMID: 7568155; PubMed Central PMCID: PMC40822.

87. Oyallon J, Vanzo N, Krzemien J, Morin-Poulard I, Vincent A, Crozatier, M. Two Independent Functions of Collier/Early B Cell Factor in the Control of Drosophila Blood Cell Homeostasis. PLoS One. 2016;11(2):e0148978. Epub 2016/02/13. doi: 10.1371/journal.pone.0148978 PONE-D-15-45333 [pii]. PubMed PMID: 26866694; PubMed Central PMCID: PMC4750865.

88. Panettieri S, Paddibhatla I, Chou J, Rajwani R, Moore R, Goncharuk T, et, al. Discovery of aspirin- triggered eicosanoid-like mediators in a Drosophila metainflammation-blood tumor model. J Cell Sci. 2019. Epub 2019/09/29. doi: 10.1242/jcs.236141. PubMed PMID: 31562189.

89. Tokusumi T, Sorrentino RP, Russell M, Ferrarese R, Govind S, Schulz, RA. Characterization of a lamellocyte transcriptional enhancer located within the misshapen gene of Drosophila melanogaster. PLoS One. 2009;4(7):e6429. Epub 2009/07/31. doi: 10.1371/journal.pone.0006429. PubMed PMID: 19641625; PubMed Central PMCID: PMC2713827.

90. Igaki T, Kanuka H, Inohara N, Sawamoto K, Nunez G, Okano H, et, al. Drob-1, a Drosophila member of the Bcl-2/CED-9 family that promotes cell death. Proc Natl Acad Sci U S A. 2000;97(2):662-7. Epub 2000/01/19. PubMed PMID: 10639136; PubMed Central PMCID: PMC15387.

91. Struhl G, Basler, K. Organizing activity of wingless protein in Drosophila. Cell. 1993;72(4):527-40. Epub 1993/02/26. doi: 0092-8674(93)90072-X [pii]. PubMed PMID: 8440019.

92. Paddibhatla I, Lee MJ, Kalamarz ME, Ferrarese R, Govind, S. Role for sumoylation in systemic inflammation and immune homeostasis in Drosophila larvae. PLoS Pathog. 2010;6(12):e1001234. Epub 2011/01/05. doi: 10.1371/journal.ppat.1001234. PubMed PMID: 21203476; PubMed Central PMCID: PMC3009591.

93. Condie JMJ.A. Mustard JA, Brower. DL. Generation of anti-Antennapedia monoclonal antibodies and Antennapedia protein expression in imaginal discs. Drosophila Information Service. 1991;70:52-4.

94. Kurucz E, Vaczi B, Markus R, Laurinyecz B, Vilmos P, Zsamboki J, et, al. Definition of Drosophila hemocyte subsets by cell-type specific antigens. Acta Biol Hung. 2007;58 Suppl:95-111. Epub 2008/02/27. doi: 10.1556/ABiol.58.2007.Suppl.8. PubMed PMID: 18297797.

95. Brower DL, Wilcox M, Piovant M, Smith RJ, Reger, LA. Related cell-surface antigens expressed with positional specificity in Drosophila imaginal discs. Proc Natl Acad Sci U S A. 1984;81(23):7485-9. Epub 1984/12/01. PubMed PMID: 6390440; PubMed Central PMCID: PMC392171.

